# Selection and transmission of the gut microbiome alone shifts mammalian behavior

**DOI:** 10.1101/2025.01.21.634013

**Authors:** Taichi A. Suzuki, Tanja Akbuğa-Schön, Jillian L. Waters, Dennis Jakob, Dai Long Vu, Mallory A. Ballinger, Sara C. Di Rienzi, Hao Chang, Ivan E. de Araujo, Alexander V. Tyakht, Ruth E. Ley

**Affiliations:** Department of Microbiome Sciences, Max Planck Institute for Biology; Tübingen, Germany; College of Health Solutions and Biodesign Institute for Health Through Microbiomes, Arizona State University; Tempe, AZ, USA; Department of Ecology and Evolutionary Biology, Cornell University; Ithaca, NY, USA; Center for Advanced Biotechnology and Medicine, Rutgers University; Piscataway, NJ, USA; Department of Neuroscience and Cell Biology, Rutgers University; Piscataway, NJ, USA; Department of Neuroscience, Icahn School of Medicine at Mount Sinai, New York, NY, USA; Department of Body-Brain Cybernetics, Max Planck Institute for Biological Cybernetics; Tübingen, Germany; Cluster of Excellence EXC 2124 Controlling Microbes to Fight Infections, University of Tübingen, Germany

## Abstract

Natural selection acts on phenotypic variation, which is influenced by genetic and environmental factors including the microbiome. Whether microbiome-mediated host phenotypes can be selected and transmitted remains untested in vertebrates. Here, we first identified locomotor activity as a trait transmissible through the gut microbiome in mice. We then performed a selection experiment, where we serially transferred microbiomes from low-activity mice to independently bred germ-free mice. Over four transfer rounds, microbiome transfer significantly reduced locomotor activity. Reduced locomotion was associated with increased Lactobacilli and their metabolite indolelactate. In a test of causality, independent administration to the mouse gut of *Lactobacillus johnsonii* or indolelactate reduced locomotion. These findings demonstrate that selection and transmission of microbes can modulate host traits independently of selection on the mammalian genome.

**One Sentence Summary:** Serial transfer of the gut microbiome of low-activity mice to germfree recipients reproducibly transferred activity levels, revealing the microbiome’s potential to impact the host’s response to selection without host genomic changes.

## Main Text

Natural selection acts on the variation in phenotypic traits present within a population of a given species. Host genetic factors, environmental factors, and their interactions, confer the variation observed between individuals. In addition, host-associated microbiomes are increasingly recognized as important sources of host phenotypic variation (*1–7*). Microbiomes influence host developmental processes and niche space, with potential impacts on their host species’ evolutionary trajectory (*1–7*). Within a single host genotype, variation in the microbiome can also produce different phenotypes. In examples of microbe-driven host plasticity, aphids with the same genotypes can exhibit different parasitoid defense (*8*), body colors (*9*), and thermotolerance (*10*), due to variability in their symbiotic bacteria alone. In such relatively simple systems, it is clear that the symbionts can shape adaptive host phenotypes (*11*). It remains to be shown whether microbiota within complex vertebrate microbiomes can similarly confer host traits that act as the basis for selection (*12*, *13*).

Selection on microbiome-driven host traits can occur if the microbiome component driving the trait is present over multiple generations of host (*1–7*). In species of insects that vertically transmit their endosymbionts with high fidelity, this condition is met (*5*). In the more complex setting of mammalian gut microbiomes, transmission routes are mixed, but there is evidence of vertical transmission in an increasing number of microbial species (*14–16*). Moreover, certain microbial species in human gut microbiomes show evidence of transmission within genetically related individuals over thousands of generations (*17*). Such long-term codiversification patterns are observed widely across primates (*18*, *19*) and other vertebrates (*20*). These observations across species provide robust evidence that vertical inheritance of specific members of the gut microbial community is widespread in vertebrates. This sets the stage for microbially-conferred host phenotypes to be passed intergenerationally and potentially selected upon, in which case phenotype-conferring microbes would act similarly to host genes.

Disentangling how the host genome and the microbial community independently respond to host-level selection is challenging and has not yet been demonstrated in animals. A powerful approach is to use artificial selection experiments (*12*, *21*). In one-sided host-microbe selection experiments (*12*), microbiomes are selected from donor hosts exhibiting a desired trait and serially transferred to independently generated germfree recipients. When performed in plants, such experiments have demonstrated that selected soil microbiomes can induce changes in host traits over time, including plant biomass (*22*), flowering time (*23*), drought tolerance (*24*), and salt tolerance (*25*). Notably, these changes in plant traits occurred relatively quickly, within five rounds of transfer. Compared to animals, plants have far fewer opportunities to transfer microbiomes to their offspring. While the results in plants show the potential to use the soil microbiome to drive desirable host traits, their relevance to the ecology and evolution of plants in natural settings may be limited.

The only equivalent animal study to date was conducted using *Drosophila melanogaster* (*26*). Microbiomes from the fastest enclosing flies were transferred into new media with sterile eggs from stock flies. However, despite a well-controlled design with replicates, the developmental time of the selection lines did not significantly differ from those of the random control lines after four rounds of transfer (*26*). Thus, in contrast to plants, there is a lack of empirical evidence supporting the potential for components of the microbiome to confer adaptive host traits in animals.

To address this gap, we performed a one-sided host-microbe selection experiment on gut microbiome-driven traits in the house mouse (*Mus musculus domesticus*). We first assessed wild-derived *Mus musculus* inbred lines as potential donors (Fig. 1A) because (i) intraspecific microbiomes from wild-derived inbred lines mitigate losses compared to interspecific fecal transfer (*27*), and (ii) a diverse wild-derived starting community is expected to improve the selection efficiency over domesticated microbiomes that have lost ancestral variation (*12*, *28–30*). We used the well-characterized and highly inbred line C57BL/6NTac as recipient germfree mice to mitigate the effects of host genetic variability on trait variation (*12*, *28*).

**Figure 1.**
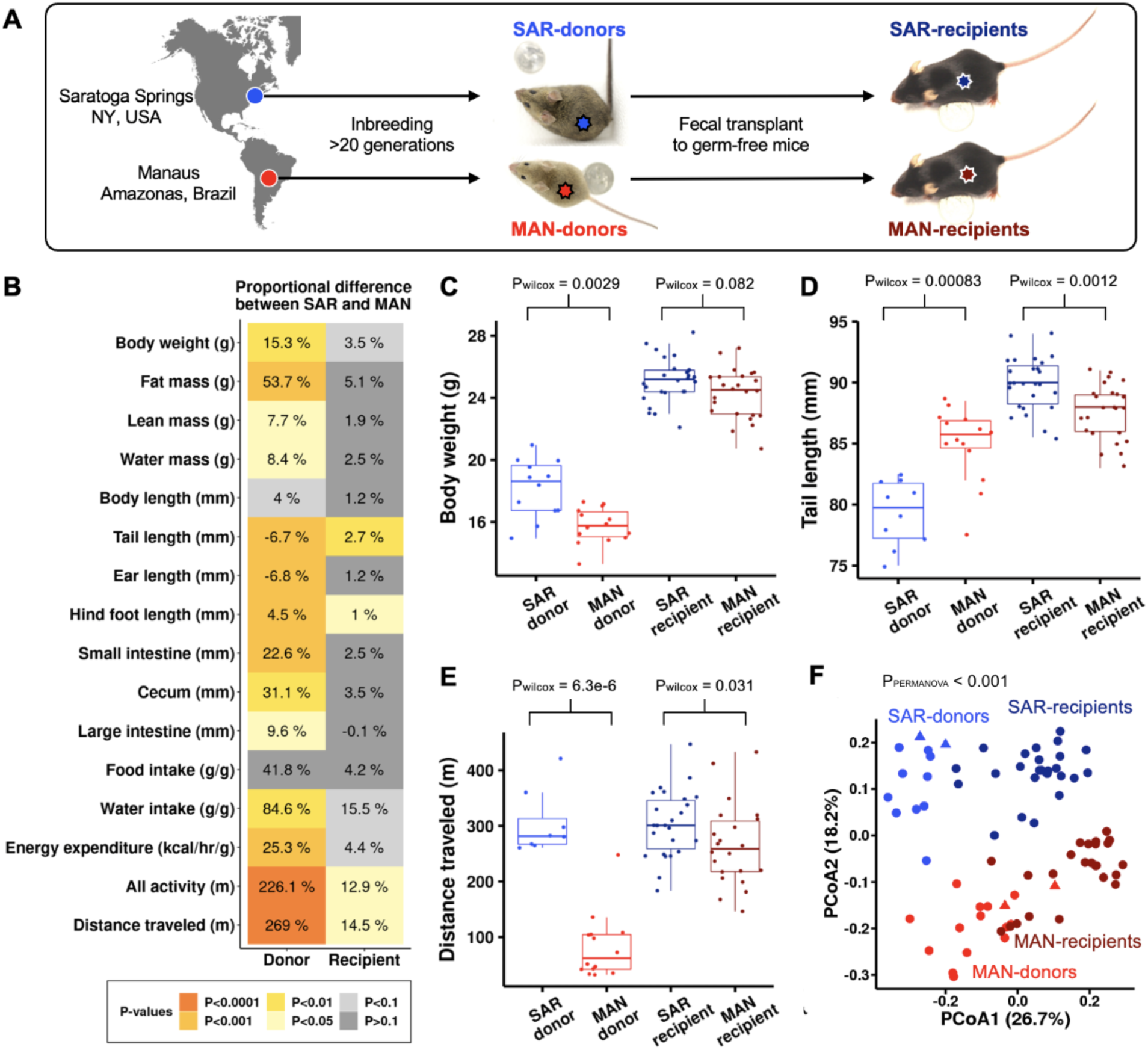
Host traits affected by microbiome. (A) Experimental design. Feces from two wild-derived inbred lines of house mice (SAR donors and MAN donors) were transferred to the cages of germfree C57BL/6NTac (SAR recipients and MAN recipients). (B) Proportional differences (%) of traits between SAR donors (n=10) and MAN donors (n=14) and between SAR recipients (n=26) and MAN recipients (n=24), all single-housed males. Positive values show greater trait values in SAR compared to MAN and negative values show the opposite. Food intake, water intake, and energy expenditure are corrected for body mass. Colors indicate Wilcoxon rank sum test uncorrected p-values. (C-E) Examples of donor and recipient traits: (C) Body weight, (D) Tail length, and (E) Distance traveled. Pwilcox: Wilcoxon rank sum test uncorrected p-values. (F) Mouse fecal microbiome between-sample diversity displayed as Principal Coordinates Analysis using Bray-Curtis distances. PERMANOVA results and the percentage of variation explained by PCoA1 and PCoA2 are indicated. Circles represent fecal microbiomes from each mouse; triangles are the fecal samples used for the transplant experiment at the start and end of the experiment.

We assessed two inbred mouse lines as microbiome donors: (i) the SAR line, originating in Saratoga, New York, USA, and (ii) the MAN line, originating in Manaus, Brazil (*31*). We characterized variation in 16 physical and behavioral traits in both lines (Fig. 1B), including locomotion-related traits measured with an automated cage system (*32*)(Methods). We observed significant differences between male SAR and MAN lines for body weight, tail length, and locomotion behavior (Fig. 1B-E; Figure S1), as previously reported (*31*, *33*, *34*). Traits with the largest mean difference for SAR versus MAN lines were ‘total activity’ (*i.e.*, sum of all movements), and ‘distance traveled’ (*i.e.*, sum of walking and running; Fig.1 B&E).

Next, we assessed the same set of traits in male C57BL/6NTac mice after conventionalization via fecal transfer from SAR and MAN donors (Fig. 1A; Methods). Overall, we observed that the transfer of microbiomes from SAR and MAN donors transmitted the differences observed in donor lines to the recipients, albeit to a greater extent for the behavioral traits compared to the morphological traits (Fig. 1B-E, Table S1, Figure S1). Only the tail length failed to phenocopy, and in fact the phenotype difference observed in the donor lines was reversed (Fig. 1D). Specifically, distance traveled showed the strongest phenocopying via microbiome transfer (Fig. 1B&E). Distance traveled significantly differed between SAR and MAN donors (Wilcoxon test uncorrected p = 0.0000063) and between SAR and MAN recipients (Wilcoxon test uncorrected p = 0.031) with the largest effect sizes within donors (269%) and within recipients (15%) (Fig. 1B&E). The differences in distance traveled remained significant after accounting for body weight at inoculation and batch effects (See Methods, Fig. S2, Likelihood ratio test p < 0.05). Our findings are consistent with quantitative genetic studies that generally show lower genetic contribution and higher environmental influence on behavioral traits compared to morphological traits (*35*).

Finally, we confirmed that the microbiomes of the SAR and MAN mice differed, and these differences in donor-mouse microbiomes could be partially transferred to recipient mice (PERMANOVA, F = 15.5, p < 0.001) (Fig. 1F, Table S2, Table S3). Based on these combined results, we chose distance traveled as the microbiome-induced trait to select for in the microbiome-transfer experiment.

## Microbiome selection drives mouse activity behavior

We set up a one-sided microbiome selection experiment (Fig. 2A; Methods) using the SAR line as microbiome donors and with the goal of reducing distance traveled. We chose to reduce (rather than increase) distance traveled as this reflects the direction of adaptation that has occurred in nature; ancestral mice from higher latitudes were introduced to lower latitudes, where activity levels are observed to be lower (*36*, *37*). We set up two lines: (i) the “selection” line, in which the two male mice with the least distance traveled after 24 hours at 5-6 weeks of age were chosen as donors for the next round, and (ii) the “control” line, in which the two mouse donors were selected at random (Fig. 2A). Recipients were inoculated at 3-4 weeks of age through coprophagy, with each round lasting 2 weeks, for 4 rounds (N0-N4; Fig 2A).

**Figure 2.**
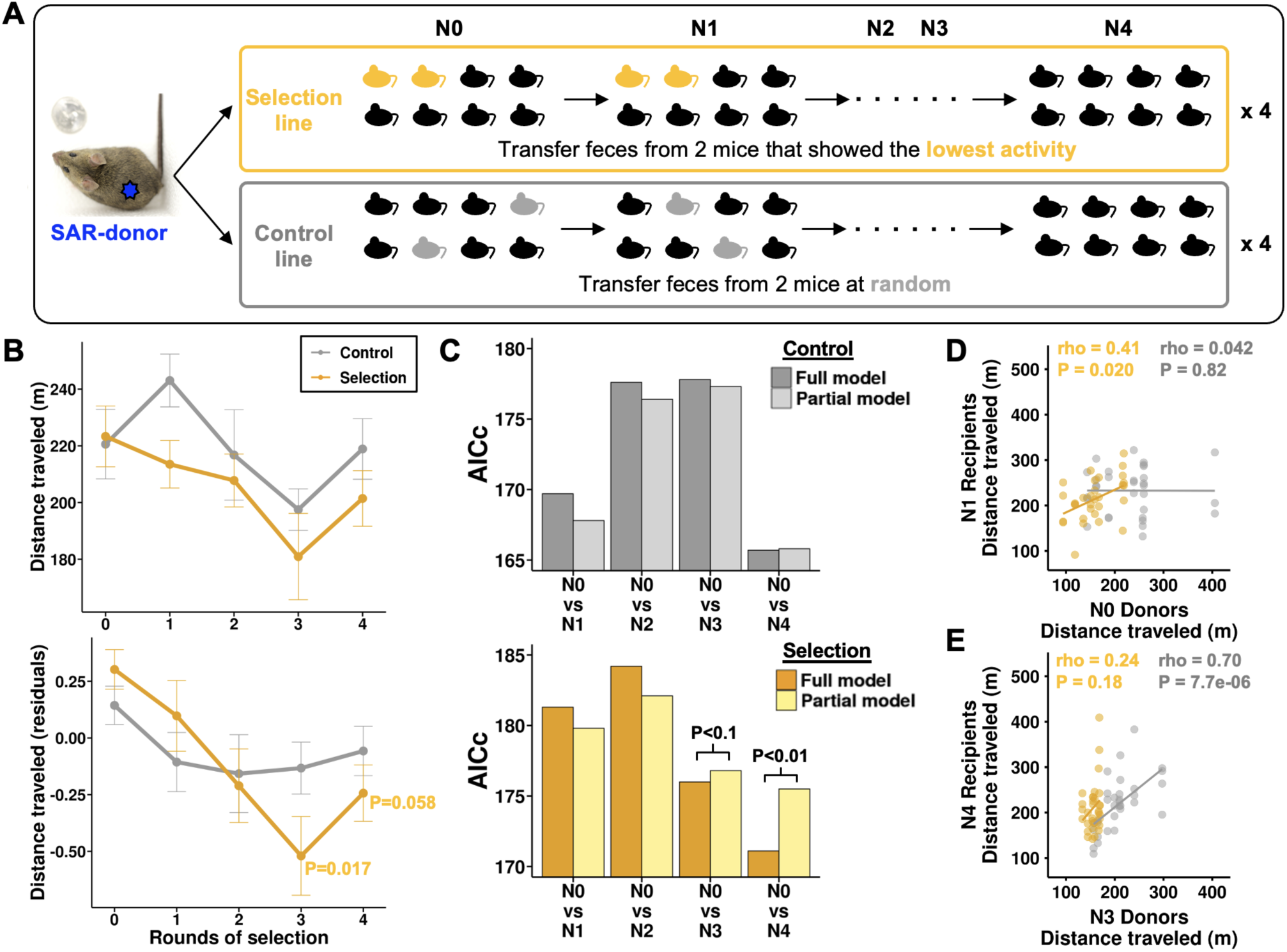
Selection and transmission of microbiome drive activity behavior. (A) Experimental design. SAR-donor mouse feces were placed in germfree recipient mouse cages. Recipients were split into two 8-mouse groups. Selection line (yellow): selection for the least distance traveled; Control line (gray): selected randomly. Process repeated four rounds (n=311 germfree mice). (B) Medians and standard errors of distance traveled for selection and control line mice. Upper panel: raw data in meters; Lower panel: residuals accounting for initial body weight and batch effects. P-values of Wilcoxon rank sum test comparing before (N0) and after selection (N1, N2, N3, and N4) are indicated: p > 0.1 are not displayed. (C) Results of model comparisons of control lines (upper panel) and selection lines (lower panel). The full model includes the distance traveled as the response variable; rounds of transfer and body weight at inoculation as fixed and batch as random effects. The Partial model excludes rounds of transfer. AICc is plotted and p-values are based on the likelihood ratio test. (D-E) Spearman’s rho correlations of distance traveled between N0 donors and N1 recipients (D) and between N3 donors and N4 recipients (E). Control line (gray); Selection line (yellow).

We observed that selection for low mouse activity behavior via transfer of the microbiome significantly reduced the median distance traveled over the whole course of the experiment (Fig. 2). As expected, in the N0 transfer time point, the control and selection lines had similar median distances traveled (p>0.05; Fig. 2B; Table S4). Correcting for starting weight and batch effects, the median distance traveled decreased over three and four rounds of transfer in the selection line (N0-N3: Wilcoxon test P = 0.017; N0-N4: Wilcoxon test P = 0.058), but not in the control line or when using the raw data (Fig. 2B). We obtained a similar result from model comparisons based on likelihood ratio tests (Fig. 2C, Table S4). These results indicate that selection and microbiome transmission were effective at reducing the distance traveled.

Next, we tested whether the distance traveled for one round of transfer was perpetuated to the next through microbiome transmission. We observed a significant positive correlation of distance traveled between the donors and recipients using all data (one round to the next; rho = 0.193, p = 0.0021, Fig. S3). Interestingly, the temporal patterns of phenocopying differed between the selection and control lines (Fig. 2D&E, Fig. S3). The selection line showed a significant phenocopying of distance traveled at the start of the experiment (N1-N2 rounds of transfer, rho = 0.41, p = 0.020, Fig. 2D), but not for the rest of the time points (Fig. S3). In contrast, the control line showed significant phenocopying of distance traveled only at the end of the experiment (N3-N4 rounds of transfer, rho = 0.70, p = 7.7e-06, Fig. 2E, Fig. S3). Overall, our results support a significant phenocopying of distance traveled through microbiome transfer, consistent with the findings from the wild-derived mice in this study (Fig. 1E) and a previous study using Diversity Outbred donors (*38*). Moreover, our observations are consistent with theoretical predictions (*21*, *39*) that selection should deplete the microbiome variation affecting distance traveled at the start of the experiment in the selection lines, while stochastic processes are more likely to maintain this variation through the end of the experiment in the control lines.

## Signatures of selection within the microbiome

Our experiments reveal that both treatment and rounds of transfer altered the gut microbiome. Analysis of the cecal microbiome diversity showed that microbiome richness (alpha diversity) was similar for the two treatments and decreased with rounds of transfer (Fig. 3A, Fig. S4; Table S5). The gut microbiomes of mice from the control and selection lines started off with similar composition (beta diversity; N0) then diverged significantly from the N0 and with each other over the rounds of transfer (Fig. 3B&C, Table S6). We detected a significant effect of treatment (selection vs. control) on Bray-Curtis dissimilarity, but the effect was relatively weak (PERMANOVA, F = 2.8, p = 0.006, Fig. 3B). In accord, rounds of transfer had a greater effect than treatment on the overall microbiome composition (PERMANOVA, F = 10.2, p < 0.001, Fig. 3B). Microbiome differences between control and selection lines were evident at the later rounds of transfer (N0-N2, PERMANOVA, p > 0.05; N3&N4, PERMANOVA, p < 0.05, Fig. 3C). Overall, the parallel shifts observed in both host traits and microbiome diversity between control and selection lines at later rounds of transfer (Fig. 2B, C, and Fig. 3C) support the notion that changes in behavior were influenced by changes in the microbiome. The observed loss of richness and the divergence between treatments over time are consistent with the stochastic loss of species caused by community bottlenecking and divergent ecosystem-level selection (*12*, *21*, *40*).

**Figure 3.**
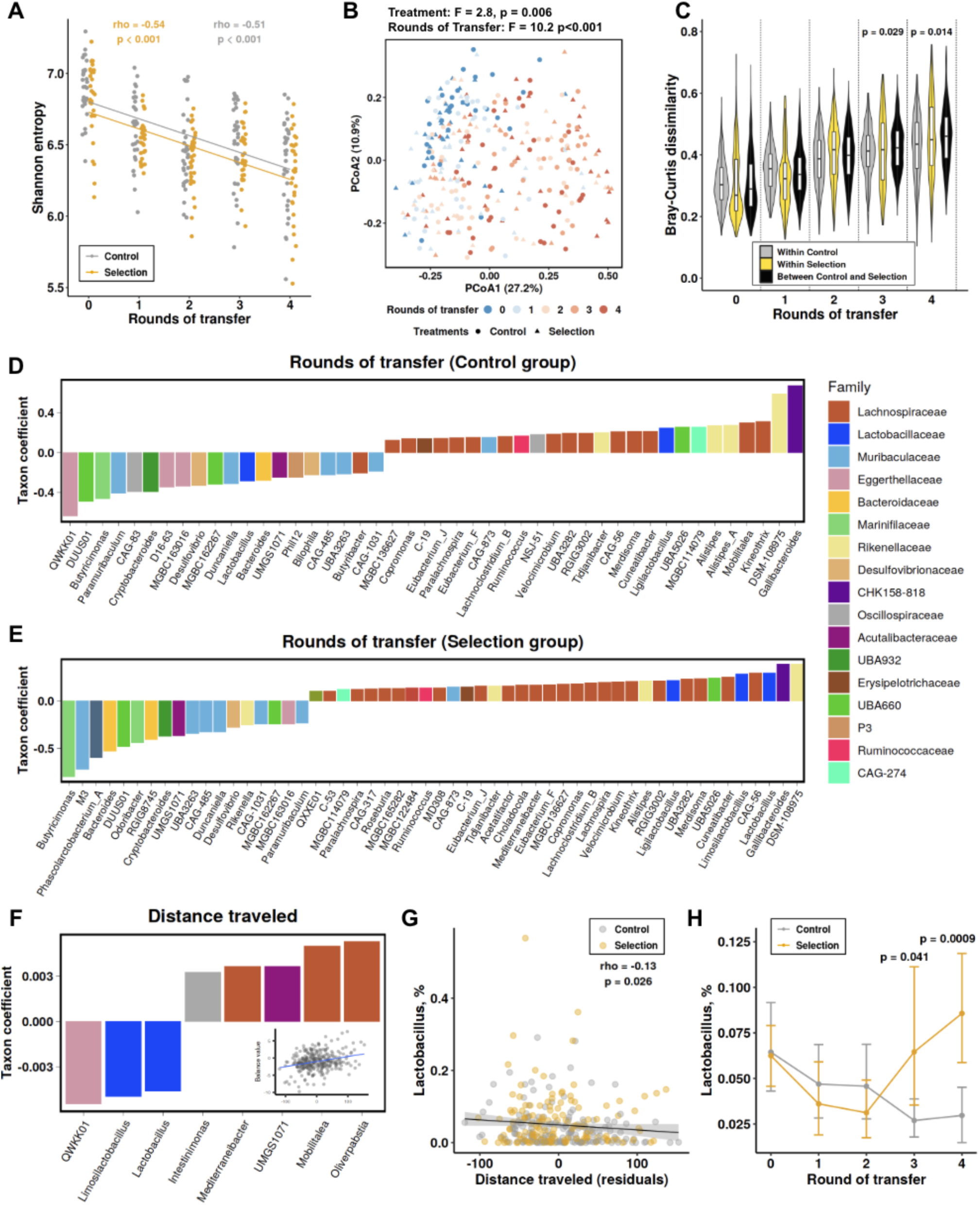
Microbiome changes in the selection experiment. (A) Shannon entropy changes across rounds of transfer, Spearman rho and p-value are shown. (B) PCoA plot of Bray-Curtis dissimilarity colored by rounds of transfer, with different shapes corresponding to treatments; PERMANOVA results are shown. (C) Bray-Curtis dissimilarity within Control lines (gray), within Selection lines (yellow), and between Control and Selection lines (black); p-values are based on pairwise PERMANOVA testing the effect of treatment on Bray-Curtis dissimilarity within each round of transfer. (D) Nearest balance (NB) associated with rounds of transfer in the Control group (n = 152), showing balance coefficients and per-sample computed values, with microbial taxa having positive values being enriched and those with negative values depleted across rounds. (E) Similar NB analysis for the Selection group (n = 151). (F) NB associated with distance traveled. (G) Correlation between distance traveled and *Lactobacillus* abundance. (H) Changes in *Lactobacillus* over time, presented as means with bootstrapped 95% CI; P-values indicate Wilcoxon tests comparing Selection and Control lines within rounds of transfer. Colors represent Selection (yellow) and Control lines (gray) in panels A, C, G, and H.

To identify specific microbial taxa whose ratios of relative abundances associated with rounds of transfer or treatment, we conducted a compositionally aware nearest balance (NB) analysis (Methods). The NB method robustly identifies associations by evaluating ratios of bacterial taxa rather than their individual abundance values (percentages) and aggregating them into a single fraction (*i.e*., balance) (*41*). We observed an increase in the balance numerator for the following taxa with rounds of transfer (*i.e*., these taxa increased; adjusted for treatment effects): *Gallibacteroides* (family CHK158-818 of the Bacteroidales), *DSM-108975* (Rikenellaceae), and many genera belonging to Lachnospiraceae and Rikenellaceae (Fig. S5 A; Table S7). On the other hand, the balance denominator taxa (decreased with rounds of transfer) included *QWKK01* (hereafter, *Enterorhabdus* as per GTDB Release 220), *Butyricimonas*, Muribaculaceae and other families. This indicates that the serial transfer alone influenced microbiome composition over rounds of transfer.

Selection for low activity resulted in an increase in *Enterorhabdus, Lactobacillus* and *Limosilactobacillus,* and four other genera, along with a reduction in proportions of *M3* (Muribaculaceae family) and eight diverse genera, when controlling for the effects of rounds of transfer (Fig. S5 B). The influence of selection on microbiome dynamics are well-illustrated through a similar balance analysis stratified by treatment (Fig. 3D,E). The described patterns were generally confirmed by the NB analysis at both the species level and the metagenome-assembled genome (MAG) level (see Methods; Tables S8-S24). The effects of selection on microbial community composition became apparent as the balance values shifted with rounds of transfer, starting with initial similarity between control and selection lines (N0) and leading to divergence in later selection rounds (Fig. S5 C). These findings suggest that the effects of selection and rounds of transfer on the microbiome were largely distinct. Yet, for overlapping taxa they strongly counteracted each other, with selection suppressing *Gallibacteroides* and *DSM-108975*, along with various Lachnospiraceae members, while enriching for *Enterorhabdus, Lactobacillus* and *Limosilactobacillus, Paramuribaculum*, and *D16-63* (Eggerthellaceae family).

## Microbial taxa associated with mouse locomotion

We computed the NB associated with distance traveled, controlling for the body weight at inoculation (see Methods, Fig. S2) and accounting for the same variables above. Distance traveled was associated with the depletion of the genera *Enterorhabdus, Lactobacillus*, and *Limosilactobacillus*, and the enrichment of *Intestinimonas* and several genera within the *Lachnospiraceae* (Fig. 3F). This is consistent with positive associations of selection with *Lactobacillus* and *Limosilactobacillus* (Fig. S5 B). Although both control and selection lines showed a significant correlation between the NB value and distance traveled, the relationship was slightly stronger in control lines compared to selection lines (Fig. S6). This aligns with the selection process diminishing variability in distance traveled within selection lines relative to control lines (Fig. 2D&E).

Further focusing on *Lactobacillus* as a key contributing taxon, we observed significant negative correlation of its relative abundance with the adjusted distance traveled across all data (Fig. 3G). When analyzing the correlation in each selection round separately, all but one round exhibited negative slopes, with significance reached in 2 of the 5 rounds (p < 0.05). The most pronounced and almost equal correlations occurred in the two final selection rounds (N3: rho = – 0.28, p = 0.029; N4: rho = –0.27, p = 0.035) (Fig. S7). Additionally, *Lactobacillus* stood out among the three top negatively contributing taxa by the increased relative abundance in the selection line compared to the control line in later rounds of selection (Fig. 3H, Fig. S8), aligning with observed changes in host behavior (Fig. 2B&C).

## Indolelactate associates with selection, locomotion, and microbiome

We measured serum concentrations at the start (N0) and end (N4) of the experiment for 12 targeted metabolites previously linked to microbiome modulation and behavioral effects in animals (*32*). Following the correction of metabolite levels and distance traveled for the body weight at inoculation, rounds of transfer significantly explained the overall variation in metabolites (R² = 0.067, p < 0.0001), whereas selection (interaction of rounds of transfer and treatment) and distance traveled did not significantly contribute (Fig. 4A). Corrected concentrations of two metabolites decreased (cortisol, corticosterone), and one increased (thyroxine), over the course of the experiment (N0-N4, p < 0.05, Fig. 4B).

**Figure 4.**
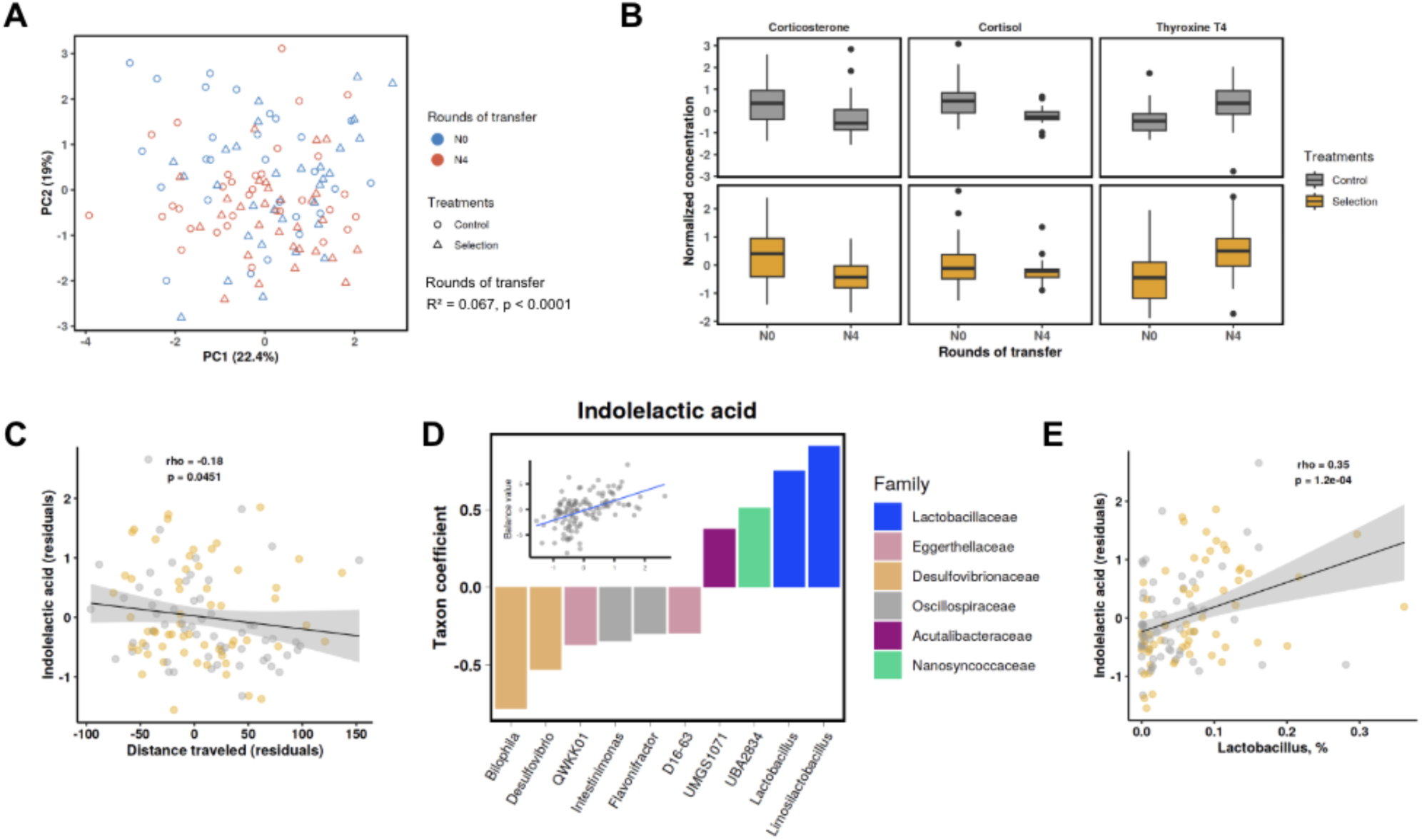
Targeted metabolite changes in the selection experiment. The levels had been corrected for the body weight at inoculation. (A) Metabolome ordination plot based on 12 metabolites, with linear mixed model results for rounds of transfer displayed. (B) Three metabolites significantly different between the start (N0) and end (N4) of the experiment. (C) Correlation between indolelactic acid and distance traveled. (D) Nearest balance associated with the indolelactic acid. (E) Correlation between indolelactic acid and *Lactobacillus*. Colors indicate Selection (yellow) and Control (gray) lines (panels B,C, and E).

Indolelactic acid (ILA) and thyroxine showed a significant negative correlation with distance traveled (Spearman rho = –0.18 and –0.19, uncorrected p = 0.0451 and 0.035, respectively; Fig. 4C and Fig. S9 A; Table S25), while for GABA it was positive (rho = 0.19, p = 0.0393; Fig. S9 B). We focused on the ILA, a microbial metabolite of tryptophan implicated in gut-brain axis communication (*42–44*) and *Lactobacillus’* capacity to catabolize tryptophan to ILA (*45*). Search for microbial genera whose ratios were associated with levels of indolelactic acid using NB yielded a balance of *Lactobacillus* and *Limosilactobacillus* (along with two other genera) to *Bilophila* and *Desulfovibrio* genera (with four others; Fig. 4D). A similar analysis at the species level revealed three closely related species in the genus *Lactobacillus* had the strongest positive association with ILA (Fig. S10 A,B). We confirmed the contribution of the leading species (*Lactobacillus johnsonii)* with a MAG-based balance analysis (Figure S9 C,D). The result is further supported by the significant positive correlation between the relative abundance of the genus *Lactobacillus* and ILA (rho = 0.35, p = 1.2E-04, Fig. 4E). These findings are consistent with the previously reported associations between *Lactobacillus* abundance, tryptophan metabolites, and hyperactivity in rodents and humans (*42–44*, *46*, *47*).

## Administration of *Lactobacillus johnsonii* and indolelactic acid to the gut reduces locomotion

Given that our NB analysis identified *L. johnsonii* as the leading and the most abundant species of the *Lactobacillus* genus associated with reduced activity, and that this species is known to produce ILA (*45*), we next tested directly the effect of *L. johnsonii* and ILA on the activity behavior of mice. First, we administered *L. johnsonii* to conventionally-raised C57BL/6J mice via oral gavage. We used an *L. johnsonii* strain LJ0 that we had previously isolated from C57BL/6J mice (*48*). Its genome encodes four genes annotated as lactate dehydrogenase. Administration of *L. johnsonii* LJ0 resulted in significantly reduced distance traveled in an open field test relative to the vehicle control treatment (Figure 5A; 2 sample t-test, p = 0.0107; 8 and 12 mice in the saline and *L. johnsonii* groups, respectively), as was speed of the mice (Figure 5B; p = 0.0136) with no changes to food intake or body weight (Figure 5 C,D; p > 0.05).

**Figure 5.**
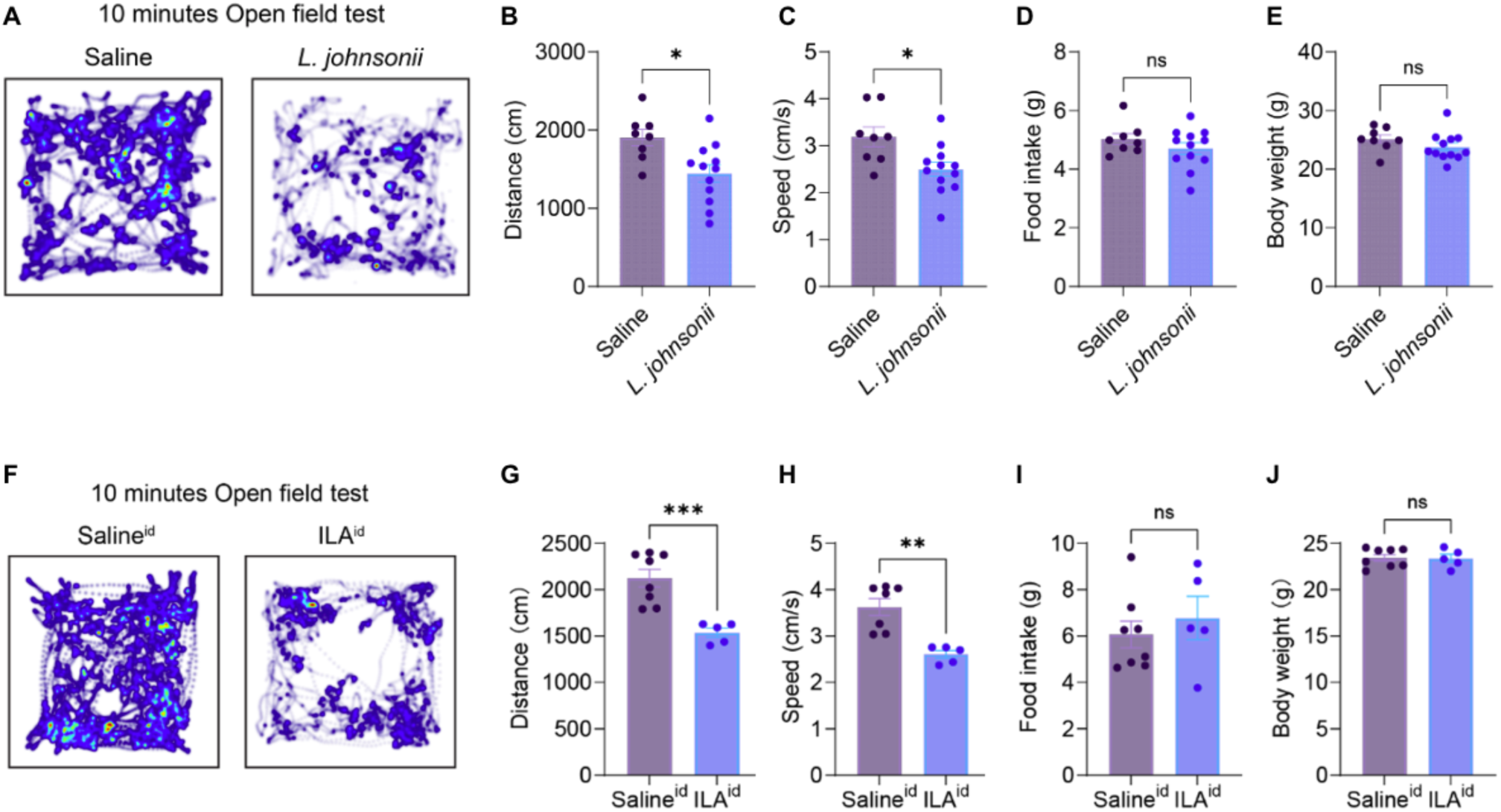
**Administration of *Lactobacillus johnsonii* or indolelactate reduces activity in mice. (**A)-(E): Open-field tests for mice with administration of saline or *L. johnsonii* solution (200 µL of ∼10^9^ cells/mL for 4 days) by oral gavage. (A) Heatmap of 10 minutes open-field tests. (B) Total distance. (C) Average speed. (D) Daily food intake. (E) Body weights. (F)-(J): same as (A)-(E) but for the mice with intraduodenal administration of saline or indolelactic acid (40 mg/kg).

To assess the effect of ILA directly in the small intestine, we administered it to the duodenum of a separate group of mice via an implanted catheter (Methods). Again, we observed a significant reduction in activity for the ILA-administered compared to the control mice (Figure 5F; p = 0.0008; 8 and 5 mice in the saline and ILA groups, respectively) and reduction in speed (Figure 5G; p = 0.0008) with no changes to food intake or weight (Figure 5H,I; p > 0.5). These results support a direct effect of ILA produced by Lactobacilli as causative in lowering the activity of mice.

## Discussion

The results of our study suggest that components of the microbiome can serve as a non-nuclear mechanism for generating host phenotypic variation that can be acted on by selection. Our study provides experimental evidence of microbiome-mediated adaptation in mammals, demonstrating that traits under selection can be transmitted through the microbiome alone.

Unlike natural populations of plants, which generally lack mechanisms for intergenerational microbial transmission except for seed-borne endophytic microbes (*49*), mammals have evolved parental care, facilitating microbial transmission from parents to offspring. Vertical transmission across generations has been observed to be widespread in different mammals (*14–16*, *50*), and can lead to host-microbial codiversification (*17*, *19*). Microbes vertically transmitted between generations and with the potential to influence outcomes of natural selection on the host are effectively acting similarly to host genes.

The microbiome-mediated adaptive plasticity demonstrated here is distinct from other transgenerational models of plasticity that require genetic variation in the host genome or epigenetic modifications such as DNA methylation (*51*). While the effect size of host trait changes in response to selection was smaller compared to previous studies in plants (*12*, *22–24*), our study supports microbiome-mediated adaptation in a vertebrate. Our results suggest that host fitness of mammals can be impacted through the acquisition of microbes without requiring genomic evolution of the host (*6*, *7*, *52*, *53*).

Pioneering studies on one-sided host-microbe selection experiments have faced criticism for their lack of control treatments, replicates, and documentation of microbiome changes (*12*). We specifically addressed these limitations in our experimental design through: (i) identifying key host traits influenced by the microbiome prior to the selection experiment, (ii) including a random control line (*12*, *26*), (iii) conducting four biological replicates per treatment, and (iv) characterizing changes to the microbiome and host metabolites. While we used coprophagy to simulate natural microbiome transmission in the mice, achieving complete fidelity of a mammalian microbiome is unrealistic in nature (*5*, *15*). Furthermore, the absence of microbiomes during the germfree recipients’ early development likely affected immune development and microbiome assembly (*54*). To address these issues, future studies should investigate the resilience of the selected microbiome by exposing animals to external microbial sources or by utilizing recipients with existing microbiomes instead of germfree mice. Including two selection lines in opposite directions (*e.g.,* high and low activity) alongside a random control line and extending the number of rounds of transfer would also enhance the statistical power to detect the effects of selection. While the host genome was kept constant in our experiment, we anticipate larger and faster changes in selected traits when both the host genome and heritable microbiome components respond to selection (*13*).

Selection on low-activity mice and transfer of their microbiomes had the effect of reducing levels of Lactobacilli in the gut and ILA in circulation. Tryptophan metabolism is increasingly implicated in the communication between the gut microbiota and central nervous system (*55*). The ILA is a bacterial metabolite of tryptophan with known immune modulatory effects through the aryl hydrocarbon receptor AhR (*56–58*). These anti-inflammatory effects extend to the central nervous system (*59*, *60*). *Lactobacillus* abundance has been linked to animal behavior in both rodents and humans (*42–44*, *61*, *62*), and to memory formation in bees via AhR (*63*). Our results suggest that the tryptophan metabolism of Lactobacilli, and subsequent ILA availability, may act on the brain to modulate activity behaviors. This provides a mechanistic hypothesis for the targets of selection within complex microbial communities (*5*).

Our work has practical applications in livestock and biomedical research (*12*). Our experimental approach demonstrates the ability to engineer culturable and unculturable microbes simultaneously *in vivo* to improve desired traits in mammals (*12*), and holds promise for uncovering novel mechanisms influencing mammalian traits and advancing microbiome-based therapeutics.

Prospectus: The gut microbiome is described as encoding a vast number of genes that extend the genomic capacity of the host. In addition, the microbiome can influence the phenotype of the host, and species within the microbiome can be passed vertically between related individuals. The results of our study indicate that specific components of the microbiome can be selected for their effect on host phenotype, independently on selection on the host genome. Thus, certain members of the gut microbiome can be viewed as non-nuclear mechanisms shaping host ecology and evolution.

## Supporting information

All Supplemental Material

## Acknowledgements

We thank Annkatrin Geysel, Matthias Neuscheler, Julia Leibßle, Silke Dauser, Heike Budde for their assistance. This work was funded by the Max Planck Society.

**Conceptualization: REL, TAS**

**Data curation: TAS, AVT, DK, DLV**

**Formal analysis: TAS, AVT, HC**

**Funding acquisition: REL, IdA**

**Investigation: TAS, AVT, HC**

**Methodology: TAS, JW, DK, DLV, SCD**

**Project administration: REL, IdA**

**Software: AVT**

**Supervision: REL, IdA**

**Visualization: TAS, AVT**

**Writing – original draft: TAS, REL, AVT**

Authors declare that they have no competing interests.

## Data and materials availability

Gut metagenomes and metagenome-assembled genomes are available for download from the European Nucleotide Archive under study accession PRJEB83173.

## Supplementary Materials

### The following includes

Materials and Methods

Supplementary Text

Figs. S1 to S9

Tables S1 to S30

References

## Material and Methods

### Animals

*Selection and transfer experiments* **-** All animal experiments were performed in accordance with the rules of the State of Baden-Württemberg, Germany, and approved by the Regierungspräsidium Tübingen (Aktenzeichen: EB 02/20M; EB 04/19M). Two wild-derived inbred lines were originally collected from Manaus, Amazonas, Brazil (MAN line) and Saratoga Springs, New York (SAR line)(*1*). SAR and MAN inbred lines were transferred from the University of California Berkeley to the Max Planck Institute for Biology, Tübingen, Germany, in the summer of 2018. The lines were maintained in Individually Ventilated Cages (IVCs) through sib-sib mating for over 20 generations at the time of this study. The animals used in this study were not rederived (*i.e.,* no embryo transfer to an existing laboratory mouse line) and the mouse lines retained population differences in the microbiome in captivity (*2*, *3*). All germ-free C57BL/6NTac mice used in this study were bred and maintained in sterile conditions (isolator bubbles and Tecniplast IsoCages P) at the Max Planck Institute for Biology, Tübingen. To ensure the germ-free status, we tested for contamination monthly by cultivation, microscopy, and/or molecular methods. The facilities were maintained at 22°C with a 12-hour light/dark cycle.

Autoclaved water and standard chow (Altromin 1314) were provided *ad libitum*. *Administrations of Lactobacilli and indolelactic acid **-*** Male C57BL/6J mice (Jax: 000664) used in the gut administration experiments were individually housed under a 12-hour light/dark cycle at the Mount Sinai animal housing facilities and fed a standard (PicoLab 5053) mouse diet. The animals were around 8-10 weeks old and weighed approximately 25-28 grams at the time of the experiments. They were used in scientific experiments for the first time. This includes no previous exposure to pharmacological agents or alternative diets. The health status of all animals was normal. All animals were individually housed for the experiments. Procedures were approved under Mount Sinai IACUC-2018-0041(de Araujo, PI).

### Fecal transplant experiments from two wild-derived inbred line donors to germ-free recipients

To characterize the traits of the MAN line and SAR line, we used a total of 14 MAN and 10 SAR mice, all males and singly housed. For recipients, we used male germfree C57BL/6NTac mice, a total of 24 MAN recipients and 26 SAR recipients, with three biological replicates (*i.e.,* each batch included 6-10 recipients per treatment). To simulate the natural transfer of microbes from parents to offspring, we utilized coprophagy. At the time of inoculation, we placed 10–15 fresh fecal pellets from 28-week-old active breeding females of each wild-derived line into the cages of newly weaned germ-free mouse recipients. Male germfree C57BL/6NTac recipient mice were weaned at 3 weeks of age, individuals were randomized from multiple litters, and two individuals were placed per cage with the feces either from the MAN female or the SAR female (MAN-recipients and SAR-recipients, respectively). We conducted this experiment in a span of seven months from the summer to the winter of 2019.

To compare the phenotypes between donors and recipients, we characterized a variety of traits related to morphology, behavior, and metabolism. Recipient’s body weights were measured every week (4, 5, 6, 7, and 8 weeks of age, n = 50) and donor body weights were measured bi-weekly (4, 6, and 8 weeks of age, n = 24) (Fig. S1). At 8 weeks of age, the body composition of live animals was measured by EchoMRI (EchoMRI LCC), including fat mass (g), lean mass (g), and total water mass (g), with a primary accumulation option of three to minimize random errors. To measure behavioral and metabolic traits, animals at 8 weeks of age inside their home cage were moved at 14:00 ± 30 mins into temperature cabinets (26°C) as previously described (*4*). Only the lid and food hoppers were replaced by Promethion cage lids (Sable Systems), while the home bedding was left intact to minimize stress on the mice. The recording started at 15:00 ± 30 mins for 24 hours. Cage locations were randomized inside the temperature cabinets for every batch. A single investigator handled all animals and cages. We measured the following traits per 24 hours; food intake (g), water intake (g), energy expenditure (mean kcal/hr using the Weir equation, indirect calorimetry based on oxygen consumption and carbon dioxide production), all activity (All Meters: sum of all movements in meters using the beam break system), and distance traveled (Ped Meters: sum of all direct locomotion in meters with a speed cut off of 1 cm/second using the beam break system). For food intake, water intake, and energy expenditure, we divided the values by body weight to account for body size differences at the time of the measurement. Gas calibration was done before every experiment and mass monitor calibration was done every four months. We used ExpeData software (Sable Systems) to extract the Promethion data using Macro13 (UMC-10.1.13-mouse.mac) and used the 24-hour measurements at five-minute resolution for all downstream analyses.

After collecting the metabolic cage measurements, 8-week-old animals were euthanized by carbon dioxide in their home cage. We collected blood samples by cardiac puncture after euthanasia. The blood was stored at room temperature for 20-30 minutes, centrifuged for 15 mins at 3000 rpm at 4°C, serum was collected, and stored at –80°C until the metabolomics analyses (see below). We also took standard morphological measurements: total length (mm), tail length (mm), hind foot length (mm), and ear length (mm). Body length (mm) was calculated as total length – tail length. The wet intestinal length was also measured: small intestine (mm), cecum (mm), and large intestine (mm). The cecum contents were stored at –80°C until DNA extraction (see below). Detailed sample information is listed in Table S26.

### Selection experiment: Serial fecal transplants from a single donor to selection and control lines

#### Selection experiment procedure

Feces from a single adult female SAR donor cage (TAS201) were placed into 16 new cages (10-15 fecal pellets each) as a starter microbial community with sterile bedding, food, and water. Newly weaned male germ-free C57BL/6NTac mice (3-4 weeks of age) were used as recipients (N0 rounds of transfer). Individuals were randomized from multiple litters, placed in cages with the starter microbiome, and separated into two groups (Selection and Control lines, 8 cages each). All animals were singly housed and the cage locations on the IVC rack were randomized. After two weeks, the recipients inside their home cage at 5-6 weeks of age were placed in temperature-controlled cabinets (set to 26°C) at 14:00 ± 30 mins. Phenotyping at 5-6 weeks and measuring at 2 PM allowed a single investigator to perform the work, thereby reducing inter-observer variability, maximizing the number of transfers within the study period, and minimizing disruption during the dark cycle. Only the lid and food hoppers were replaced by Promethion cage lids (Sable Systems) and the recording started at 15:00 ± 30 mins for 24 hours. Cage locations were randomized inside the temperature cabinets. A single investigator handled all animals and cages. After the recording, all animals (N0 rounds of transfer) were euthanized and measurements were taken (see below). For the selection line, feces from two individuals that showed the lowest distance traveled (Ped Meters) were collected. Feces from these individuals were placed into 4 new cages (10-15 fecal pellets each) with sterile bedding, food, and water (8 cages total for the selection line). For the Control line, the same procedure was performed as the Selection line, but feces from two random individuals (using an online random number generator) were collected instead (8 cages total for the control line). Within an hour of placing the feces into the new cages, newly weaned germ-free recipients (3-4 weeks of age) were placed in the cages, and inoculation occurred via coprophagy (N1 rounds of transfer). We repeated this selection procedure four times (N0 – N4 rounds of transfer). Finally, we repeated the entire procedure in parallel four times resulting in a total of four biological replicates involving a total of 311 germ-free recipient mice from summer to winter 2020.

Promethion data (*i.e.,* food intake, water intake, energy expenditure, all activity, and distance traveled), body weight, body length, tail length, and cecum were collected after two weeks post inoculation (5-6 weeks of age) from all animals involved in the selection experiment. For the first (N0) and last (N4) rounds of transfer, body compositions of live animals were measured at 5-6 weeks of age by EchoMRI (EchoMRI LCC) and serum was collected using cardiac puncture after euthanasia for metabolomics (see below). As described above, gas calibration was done before every experiment, and mass monitor calibration was done every four months. We used ExpeData software (Sable Systems) to extract the Promethion data using Macro13 (UMC-10.1.13-mouse.mac) and used the 24-hour measurements at five-minute resolution for all downstream analyses. Detailed sample information is listed in Table S27.

### Bacterial culturing, oral administration, and open field tests

*Lactobacillus johnsonii* strain LJ0 was previously isolated on MRS agar plates from the small intestine of a specific pathogen-free housed C57BL/6 mouse (*5*). The annotated genome is available in Genbank (assembly ID: GCA_002156645.1). *L. johnsonii* was suspended in sterile water and spread onto MRS agar plates (de Man, Rogosa, Sharpe, Millipore, #110660) containing anaerobic cultivation packs (Thermo Scientific, AnaeroPack™) and incubated at 37°C. The following day, bacterial cells were scraped off from the plates and were harvested by centrifugation at 2500 × g for 5 minutes and diluted in sterile saline to achieve a concentration of approximately 10^9^ cells/mL. Animals were administered either a saline solution or *Lactobacillus johnsonii* suspension (200 µL) via oral gavage daily for four consecutive days. On the fourth day, after an interval of approximately 4 hours, 10-minutes open field tests were performed separately for each individual mouse. Specifically, animals were placed in a novel Plexiglas arena (Med Associates, 25 cm × 25 cm), where a 150-W lamp positioned above the central subarea was activated to elicit natural aversion to the area, following standard protocol. Animals were tested once in this arena, and average velocity and total distance traveled were calculated using automated video analysis software (EthoVision XT 11.5, Noldus).

### Intra-duodenum catheterization, indolelactic acid (ILA) infusions, and open field tests

At the time of experimentation, animals were 8 weeks old. The abdomen and dorsal back skin of each animal were shaved and cleansed. A small incision was made on the dorsal neck, and a midline incision was created on the abdomen to exteriorize the duodenum. A purse-string suture was placed on the duodenal wall, 1 cm proximal to the start of the duodenum, through which the tip of MicroRenathane tubing (0.025”, Braintree Scientific, Braintree, MA) was inserted. The purse-string suture was tightened around the tubing, which was then tunneled subcutaneously to the dorsum through a small opening in the abdominal muscle. A small incision was made between the shoulder blades to allow for catheter exteriorization. The abdominal incision was closed using sterile sutures. Saline or ILA solution (40 mg/kg) was infused into the mice after complete recovery. Following a four-day infusion period, both control and ILA-treated animals were subjected to an open field test (described above).

### Targeted metabolomics

We selected twelve metabolites reported to affect host behavior through gut microbiome modulation (*4*) and measured their concentrations in serum from animals at the start (N0) and end (N4) of the selection experiment using LC-MS analysis performed on a HPLC system (Dionex UltiMate 3000, Thermo Fisher, USA) coupled with a high-resolution mass spectrometer (Impact II, Bruker, Germany). We used the protocol described in (*4*) for corticosterone, glutamic acid, kynurenic acid and γ-aminobutyric acid (GABA). For the other eight compounds – indole-3-lactic acid (ILA), indoxyl sulfate, indole-3-propionic acid, cortisol, tryptophan, thyroxine (T4), kynurenine and serotonin – an updated LC-MS/MS method was used with the following separation condition: stationary phase C18 Kinetex 2.6 µm; 100 A; 150×2.1 mm kept at 40°C during the analysis. Mobile phase A was water and mobile phase B was acetonitrile, both with the addition of formic acid (0.1%). The following gradient: A/B 99/1 (0 min), 60/40 (10 min), 5/95 (12 to 12.5 min), 99/1 (13 to 15 min) was applied at a constant flow rate of 0.4 ml/min. The injection volume was 5 µl. Kynurenine and serotonin were analyzed in positive ionization while the others were acquired in negative mode. Collision energy was from 15 to 35 eV.

### Generation and processing of metagenomes

DNA was extracted from frozen cecal samples using the PowerSoil DNA isolation kit (Qiagen, Valencia, CA, USA) according to the manufacturer’s protocol. We prepared metagenomic libraries as described (*6*) with slight modifications. Briefly, 1 ng of purified gDNA was used in a Nextera (Illumina, San Diego, USA) Tn5 tagmentation reaction to fragment and ligate adaptors in a single reaction, followed by a 14-cycle PCR to add sample-specific barcodes. Libraries were purified, pooled, and quantified. Size selection (400-700 bp) was performed on a BluePippin (Sage Science, Beverly, USA). Libraries were concentrated and further purified as needed using DNA Clean & Concentrator-5 (Zymo Research, Irvine, USA). Sequencing was conducted on a HiSeq 3000 System (Illumina, San Diago, USA) with 150 paired-end sequencing. We used a quality control pipeline described in (*6*). Briefly, adapter trimming and quality control filtering were conducted using Skewer 0.2.2 and bbtools “bbduk” command. Reads mapping to the human genome (GRCh37/hg19) and mouse genome (GRCm39) were filtered using bbtools “bbmap” command. The read quality was assessed using Fastqc 0.11.7 and multiQC 1.5a.

### Microbiome analyses

#### Taxonomic and functional profiling

Functional and taxonomic profiling was conducted using an in house pipeline. Briefly, taxonomic profiling was based on Kraken2 (*7*) with default parameters, and Bracken v2.2 (*8*) parameters set to “-t 10 –l S”. Functional profiling was based on HUMANn3 v3.0.0.alpha.3 (201901) (*9*). Custom databases were created using Struo2 (*10*) based on GTDB release 207 (*11*). The custom database used here is available at (http://ftp.tue.mpg.de/ebio/projects/struo2/GTDB_release207/).

During the initial component-based microbiome data analysis, we rarefied at 150,000 reads, which excluded one sample (R3N5T290) from all downstream analyses. Relative abundances, alpha diversity, and beta diversity were calculated by Qiime2 (*12*). Observed features, Shannon index, and Faith’s PD were calculated for alpha diversity. Bray-Curtis dissimilarity, unweighted– and weighted-UniFrac distances were calculated for beta diversity. Rarefied reads were also used to calculate unstratified and stratified pathways using HUMANn3 v3.0.0.alpha.3 (201901) (*9*). To test differences in beta-diversity by groups, we applied PERMANOVA in QIIME2 (*12*) using the “diversity beta-group-significance” command. To test differences in alpha diversity by groups, we applied the Wilcoxon rank sum test.

For validation purposes, we employed an alternative approach to taxonomic profiling – one based on metagenome-assembled genomes (MAGs). Due to the moderate coverage of the non-rarefied cecal metagenomes, to increase the quality of the assembly they were pooled: for the Experiment 1 – by origin (SAR or MAN) × (donor or recipient) – into 4 metagenomes, for the Experiment 2 – by replicate × round of transfer × treatment (except for all N0 round samples that were pooled into one and the donor’s sample that was not pooled) – into 34 metagenomes; yielding 38 pooled metagenomes in total. The metagenomic assemblies, binning into metagenome-assembled genomes (MAGs), taxonomic classification and dereplication into species-representative genomes (SRGs) were performed as described previously (*13*), with the MAG quality assessed using CheckM2 (*14*). Out of 127 species initially represented by the MAGs, 104 were assigned an SRG as a result of the dereplication; for 11 of the remaining ones that were taxonomically classified at the species level, their representative genomes were downloaded from GTDB. The resulting set of genomes was transformed into a custom reference database for KrakenUniq software (*15*) that was subsequently used to obtain a taxonomic profile for each non-rarefied metagenome.

#### Compositionally aware statistical analysis of microbiome

For the 310 non-rarefied metagenomes in the selection experiment, the genera with a prevalence <30% at >0.005% relative abundance were discarded, leaving 158 features (the respective number during the species level analysis was 687). The samples outlying (outside of median ± 3 sd) by body weight at inoculation (BWi) or distance traveled were discarded, leaving 303 samples. For the analyses involving metabolomic data, three samples outlying by their metabolomes were also excluded from consideration (leaving 118 samples with both metagenomes and metabolomes). After zero imputation based on Bayesian-multiplicative replacement, per-taxon read counts were CLR-transformed. The distance traveled and scaled serum concentration of each metabolite were adjusted for BWi by collecting the residuals from the respective LOESS model (ɑ = 0.75). Compositionality-aware evaluation of the association of each factor of interest to the variation of microbiome composition was carried out by applying PERMANOVA (adonis2 from vegan) to the Aitchison distance matrix using the formula: *beta-diversity ∼ distance traveled + BWi + round of transfer + treatment + round of transfer: treatment + replicate* (9999 permutations) (Table S28). For each significantly associated factor, a compositionality-aware identification of its associated taxa was performed using the Nearest Balance method (*16*). The method processes the outputs of a linear model of the normalized taxa abundance values to yield the optimal ratio (balance) of two subsets of taxa (numerator and denominator) associated with the factor. The following formulas were used in the model for obtaining the taxa coefficients for each factor of interest, with clr-transformed abundance values as a response:

*∼ adj. distance traveled + BWi + round of transfer * treatment* – for the distance traveled, round of transfer and interaction of treatment with rounds; *∼ adj. metabolite level + BWi + round of transfer * treatment* – for the levels of each metabolite, respectively. The consensus nearest balance was calculated using 100 iterations of cross-validation (train set proportion: 0.67) to include the taxa with reproducibility >80%. Color palettes from the ggsci R package were used for visualization.

### Targeted metabolome analysis

For each metabolite of interest, its body weight-adjusted scaled value was the input of a linear model (lmer function from lmerTest package) with the following formula: ∼ *round of transfer * treatment*; the resulting p-values were FDR-adjusted and the findings with p.adj < 0.1 were reported as significant.

### Statistics

#### Initial measurements with MAN and SAR donors

To test differences in trait values between groups (*i.e.,* MAN-donors vs SAR-donors, MAN-recipients vs SAR-recipients), we applied the Wilcoxon rank sum test on raw data. However, we observed significant effects of body weight at inoculation (3 weeks old) and at measurement (8 weeks old) on distance traveled at 8 weeks old, with a stronger correlation for body weight at inoculation (Fig. S2). Given that each batch of newly weaned germ-free mice included multiple litters at slightly different ages, we accounted for both batch and body weight at inoculation. To test whether the differences between groups persist after accounting for body weight at inoculation and batch effects, we created hierarchically nested linear mixed-effects models using “lmer” function in “lme4” R package. Then we conducted model comparisons using likelihood ratio tests using “lrtest” function in “lmtest” R package. We compared two nested models: (model 1) a full model, including a trait of interest as the response variable,

groups (*i.e.,* MAN-donors vs SAR-donors or MAN-recipients vs SAR-recipients), and body weight at inoculation (3 weeks of age) as fixed effects, and the batch as random effects and (model 2) a partial model, including all the same variables as the full model except excluding the group variable. The input variables were log-transformed and standardized. Both uncorrected and FDR-corrected p-values are reported (Table S1).

#### Selection experiment

To test whether distance traveled significantly differed between treatments (Control vs Selection) or between before and after selection (e.g. N0 vs N4), we applied the Wilcoxon rank sum test on raw data and residual values accounting for covariates (see below). Similar to the pilot experiment above, we compared two nested linear mixed-effects models: (1) a full model, including distance traveled as the response variable, rounds of transfer (N0 vs N1, N0 vs N2, etc.) and body weight at inoculation (3 weeks of age) as fixed effects, and the batch as random effects, (2) a partial model including the same variables as the full model except excluding the round of selection variable. The variables were log-transformed and standardized. AICc (corrected Akaike information criterion) of the models is reported. We compared the models using likelihood ratio tests using “lrtest” function in “lmtest” R package. Both uncorrected and FDR-corrected p-values are reported (Table S4).

### Supplementary Text

#### Rationale for selection experiment study design

The design of the experiment was inspired by a One-Sided Host-Microbiome Selection experiment (*17*), where the microbiome is allowed to change in response to selection on a desired trait while the host genome remains constant. For the starting microbial community, we used feces from a single SAR donor. Wild mouse-derived microbiomes are expected to enhance the translatability of laboratory mouse studies to humans (*18*, *19*). Additionally, they are expected to respond more rapidly to selection, as established communities are likely to exhibit minimal initial changes in a new recipient compared to mixed or synthetic microbial starter communities (*17*).

We chose distance traveled (Ped Meters) as the trait to select for, and in the direction to lower the trait value, for the following reasons. First, activity behaviors showed the strongest evidence of phenocopying among all traits tested, where differences in distance traveled in recipient lab mice reflected that of the donor mice (see Main Text Fig. 1). Second, we selected the high-latitude starter microbiome in the direction from high activity to low activity to reflect the direction of adaptation that occurred in nature, where a higher metabolic rate is likely more beneficial in higher latitudes, and vice versa (*20*). Although selecting for low activity has a lower bound (as opposed to selecting for high activity), the pilot experiment indicated that selection could be more effective in reducing the activity than increasing it by considering the natural activity range of the two donor mice (Main Text Fig. 1E).

While selecting the trait in both directions would likely maximize the chance to detect the changes in activity, the lack of a control line that changes stochastically has been criticized in previous studies (*17*, *21*, *22*). Thus, we decided to characterize the null changes in activity over time (Control line) to test whether the activity changes observed in the Selection line significantly differed from that of the Control line as in (*21*, *22*).

#### Microbiome diversity analyses

To confirm whether we successfully transferred the microbiomes from donors to recipients through coprophagy, we created a beta-diversity PCoA plot based on Bray-Curtis dissimilarity (Main Text Fig. 1F). The PCoA1 axis separates donors and recipients and the PCoA2 axis separates SAR and MAN microbiomes (PERMANOVA, F = 15.5, p < 0.001) supporting a successful microbiome transfer. Unweighted– and weighted-UniFrac distances also showed similar patterns (Table S3). For the three alpha-diversity metrics tested, none differed between SAR and MAN donors (Table S2), and only the Shannon index was significantly reduced in recipients compared to donors (Wilcoxon test p < 0.025), indicating a shift in evenness of the community. Together, the results support the idea that population differences in the microbiome can mediate population differences in host traits, especially activity behaviors (Main Text Fig. 1).

#### Microbiome metabolic pathway-related analysis

We also identified microbial pathways from the shotgun metagenomic data that were significantly associated with distance traveled during the selection experiment (Table S29&S30). Among 184 unstratified pathways tested, only “gluconeogenesis_III” significantly correlated with distance traveled using the cut-off of q < 0.25 accounting for body weight at inoculation and batch effects (Table S29). Among the top 10 pathways with the highest coefficients with distance traveled, five pathways are directly involved in cellular respiration ranging from glycolysis, pyruvate oxidation, and TCA cycle (*i.e.*, “gluconeogenesis_III”, “pyruvate fermentation to propanoate I”, “pyruvate fermentation to acetone”, “acetyl-CoA fermentation to butanoate II”, and “incomplete reductive TCA cycle”). All five pathways are positively correlated with distance traveled. Among the 829 stratified pathways tested, two pathways remained significant at the cut-off of q < 0.25 (Table S30). The top pathway was phosphopantothenate biosynthesis I (*Eubacterium* sp.) involved in the biosynthesis of coenzyme A, which is consistent with the unstratified results related to metabolic reactions involved in cellular respiration. The second pathway was dTDP-L-rhamnose biosynthesis I (1XD8.76 sp.) involved in the biosynthesis of lipopolysaccharides. Stratified pathways linked to *Lactobacillus* or *Limosilactobacillus* were not detected.

#### Adjustment of factors for body weight at inoculation for the microbiome and metabolomic analysis

Examination of the relationship between metabolite levels and body weight at inoculation revealed non-monotonous patterns. Because of their deviations from linear behavior, LOESS (locally estimated/weighted scatterplot smoothing) was preferred to the linear interpolation. For the distance traveled, the deviation was moderate, occurring at the few higher values, therefore, in the non-microbiome/metabolome part a linear mixed effect model was accepted for its adjustment.

#### Ecological context

The findings in this study align with previous studies in wild-derived mouse lines, where mice from colder regions display increased wheel running and nest building behaviors compared to those from warmer regions (*20*, *23*). Given the known role of higher physical activity on regulating metabolic responses to cold (*24*), these findings provide further evidence for microbiome-mediated adaptive host plasticity associated with thermoregulatory adaptation (*25*–*28*). The results suggest that both population differences in the microbiome and host genetics may contribute to maintaining adaptive behavioral variation in wild house mouse populations.

Among the morphological traits, the only significant difference observed in the recipients due to differences in the microbiome was the tail length. Interestingly, a previous study using the same wild-derived inbred lines demonstrated that tail length showed strong plasticity in response to temperature differences, while body weight showed limited plasticity (*29*). This suggests that tail length is more responsive to changes in temperature and the microbiome compared to body weight. The contrasting patterns of tail length between the recipients and donors can be attributed to differences in the microbiome’s energy extraction capabilities (*3*, *30*, *31*). The composition and function of the gut microbiome are associated with mouse body mass in the field and captivity: the SAR donors were found to produce higher levels of short-chain fatty acids compared to the MAN donors under the same environmental conditions, with no difference in food intake (*3*). This explains the trend where SAR recipients exhibited greater body weight and longer tail length compared to MAN recipients. The significant opposite patterns of phenocopying for tail length suggest that host genotypic differences primarily determine the allocation of energy to tail growth, with the microbiome playing a secondary role.

## Supplemental figures

**Figure S1.**
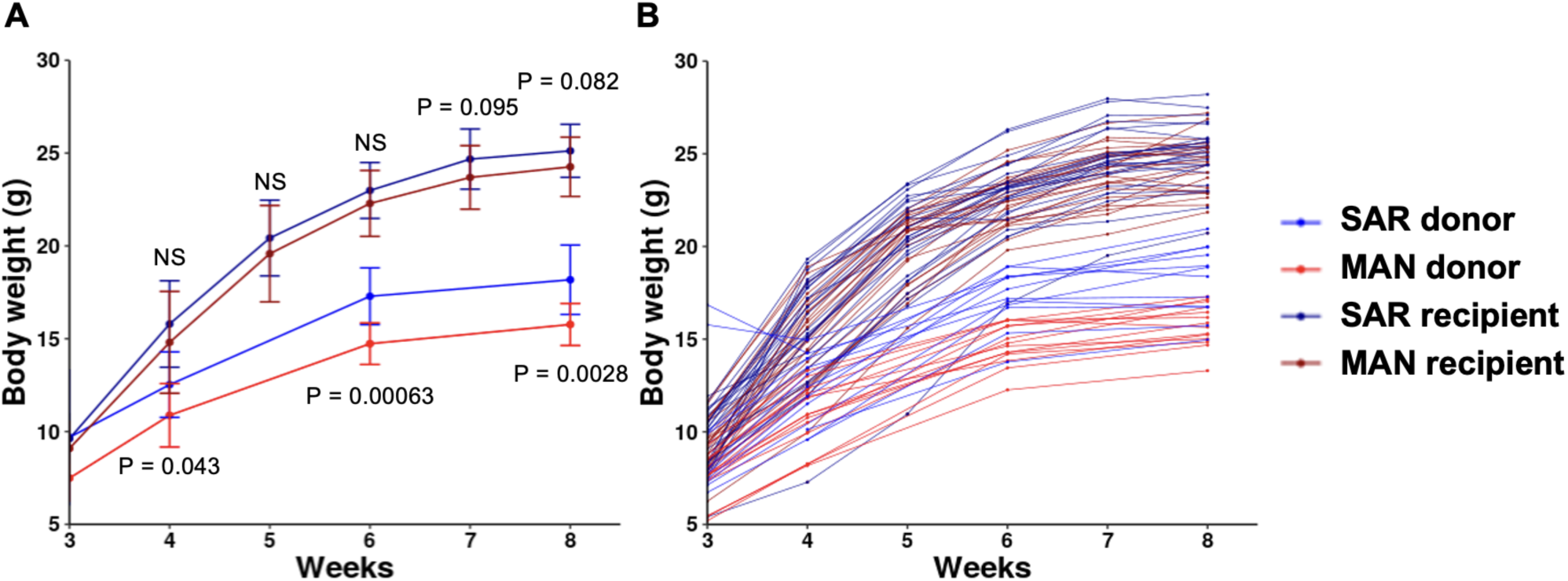
Growth curves of donors and recipients. (A) Mean body weights of SAR donors (blue), MAN donors (red), SAR recipients (dark blue), and MAN recipients (dark red). Error bars are standard deviations. All animals are male. Wilcoxon rank sum test p-values are plotted for each time point within donors and within recipients: NS p > 0.1. (B) Individual data points for body weight are plotted.

**Figure S2.**
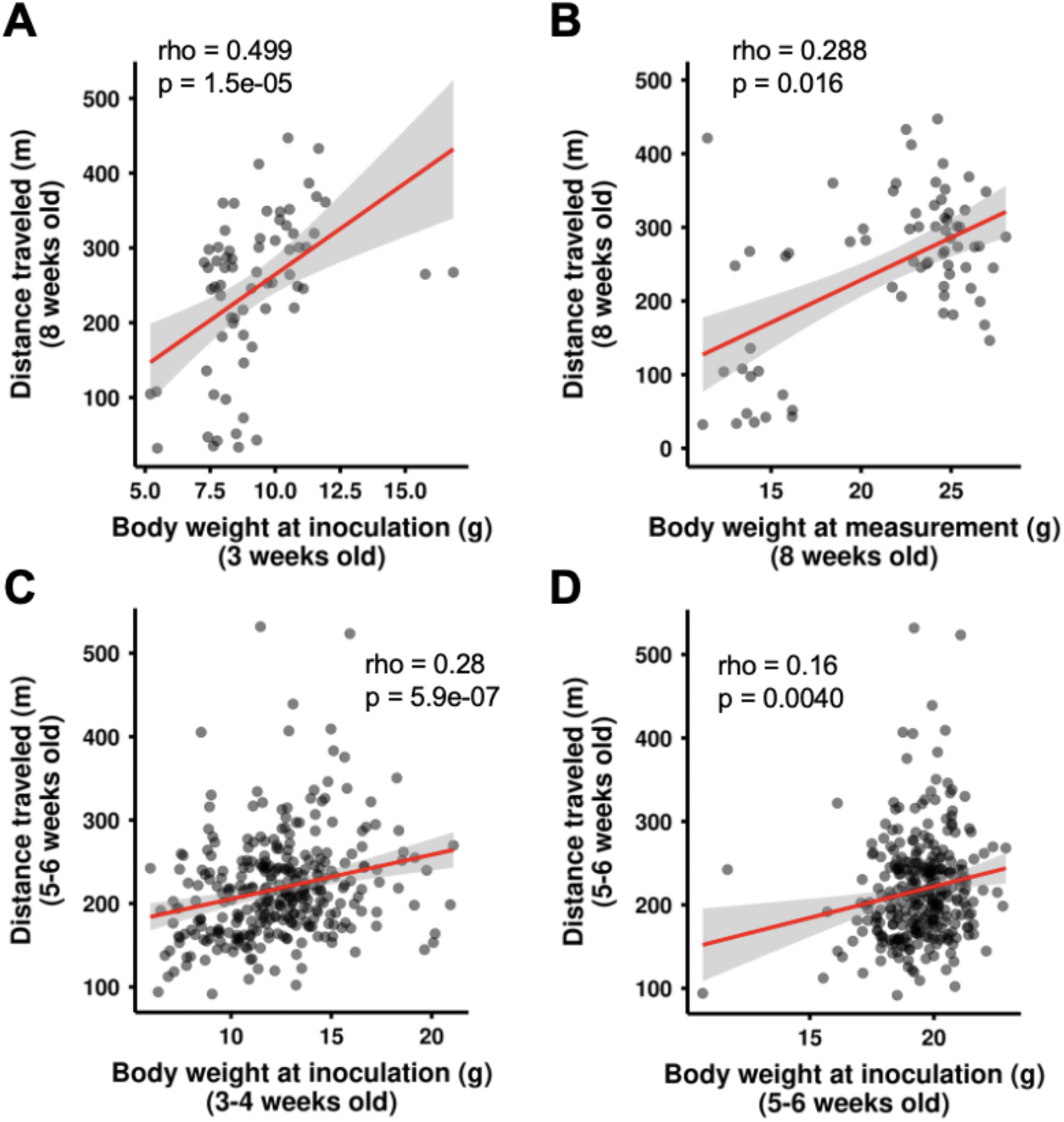
Correlations between body weight and distance traveled. Correlations (A) between body weight at inoculation (3 weeks old) and distance traveled (8 weeks old), (B) between body weight at measurement (8 weeks old) and distance traveled (8 weeks old) from the pilot experiment. Correlations (C) between body weight at inoculation (3-4 weeks old) and distance traveled (5-6 weeks old), (B) between body weight at measurement (5-6 weeks old) and distance traveled (5-6 weeks old) from the selection experiment.

**Figure S3.**
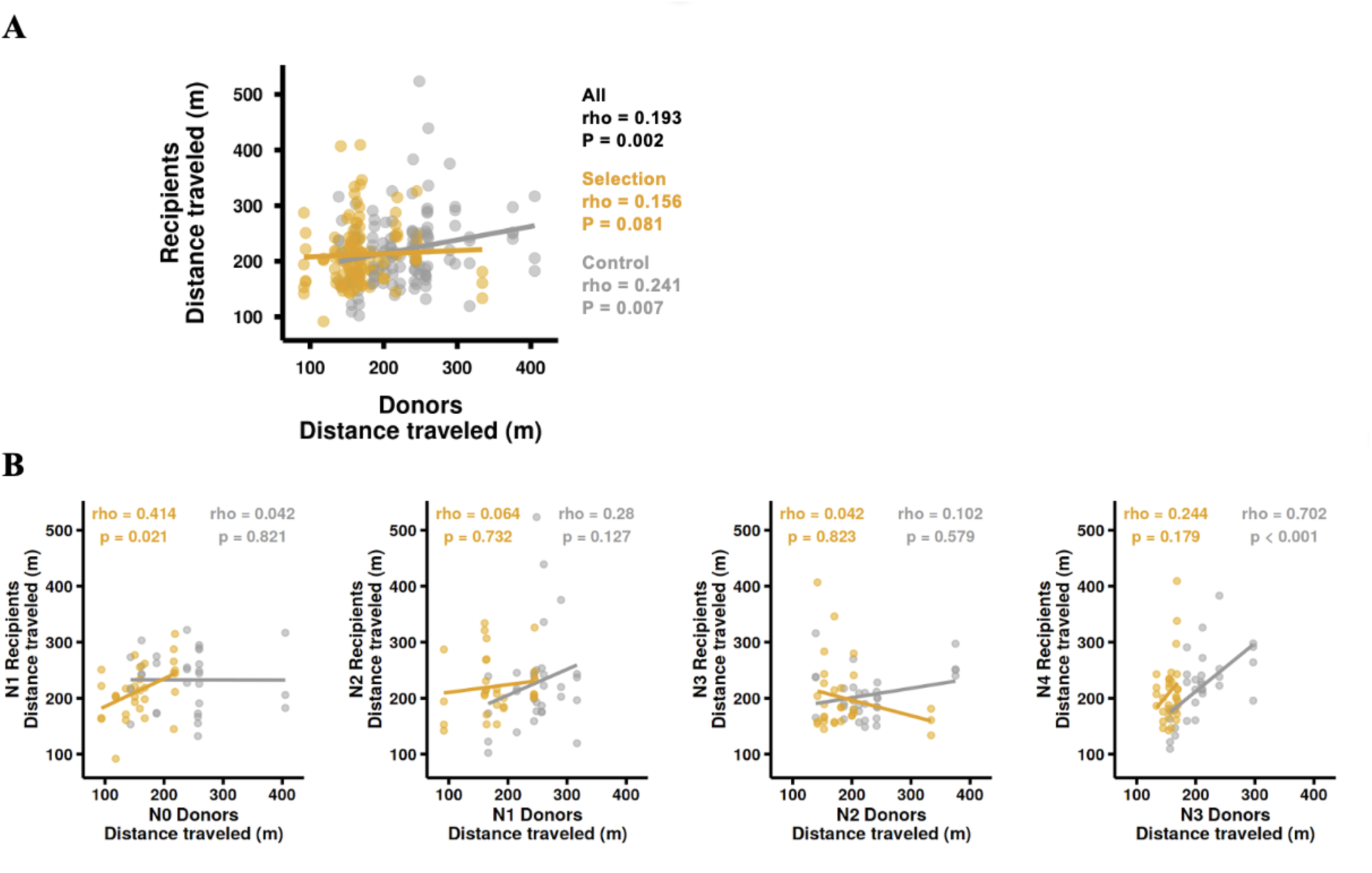
Community-level heritability of distance traveled. (A) Correlation of distance traveled between all donors and all recipients. Spearman rho and p-values are shown for all data points (black), Selection line (yellow) and Control line (gray) (B) Correlation of distance traveled between donors (N-1) and recipients (N) by each round of selection.

**Fig. S4.**
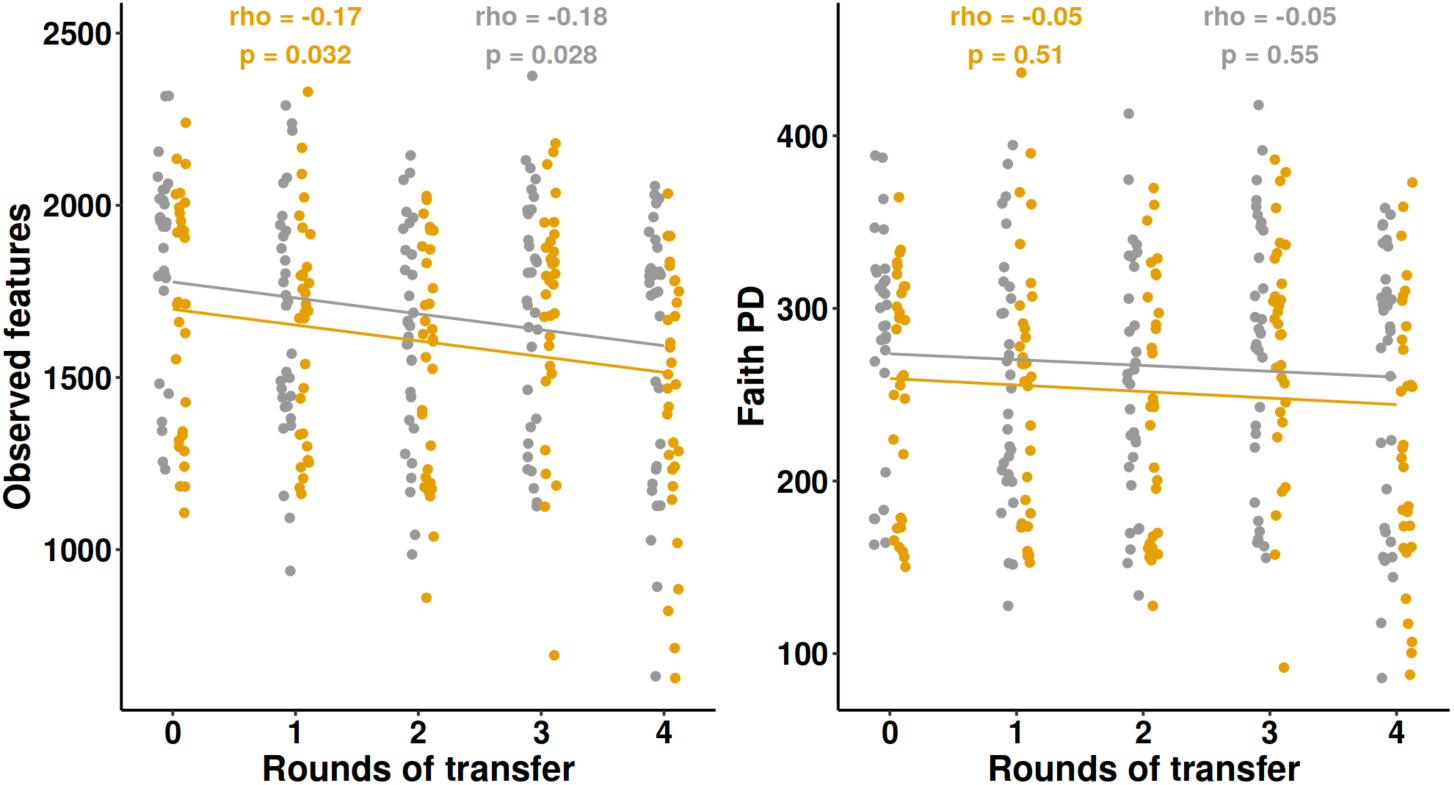
Alpha-diversity changes across time. Observed features showed a significant negative correlation with rounds of transfer. Faith PD did not significantly correlate with rounds of transfer. Colors indicate the selection line (yellow) and control line (gray). Spearman rho and p-value are shown.

**Figure S5.**
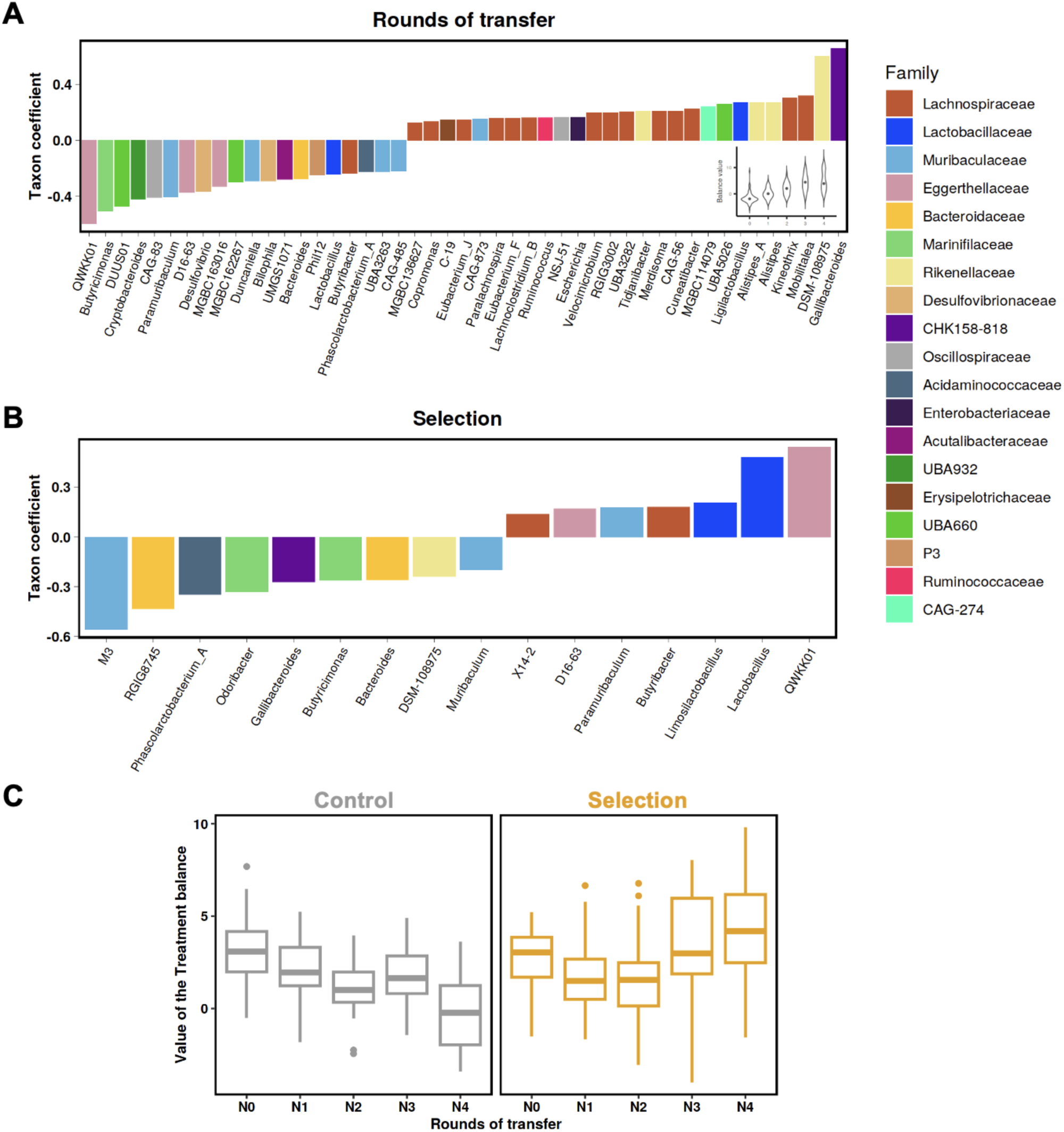
Nearest balances (NB) associated with the rounds of transfer and selection. (A) NB associated with rounds of transfer across both Control and Selection group: balance coefficients and its per-sample computed values. Microbial taxa with positive coefficients are enriched in later rounds of selection. (B) NB associated with selection (modeled as interaction of rounds of transfer and treatment). Microbial taxa with positive coefficients are enriched in the Selection lines, while those with negative coefficients are enriched in the Control lines across the rounds. (C) Dynamics of the selection-associated NB values for Control (gray) and Selection lines (yellow) across the rounds of transfer.

**Figure S6.**
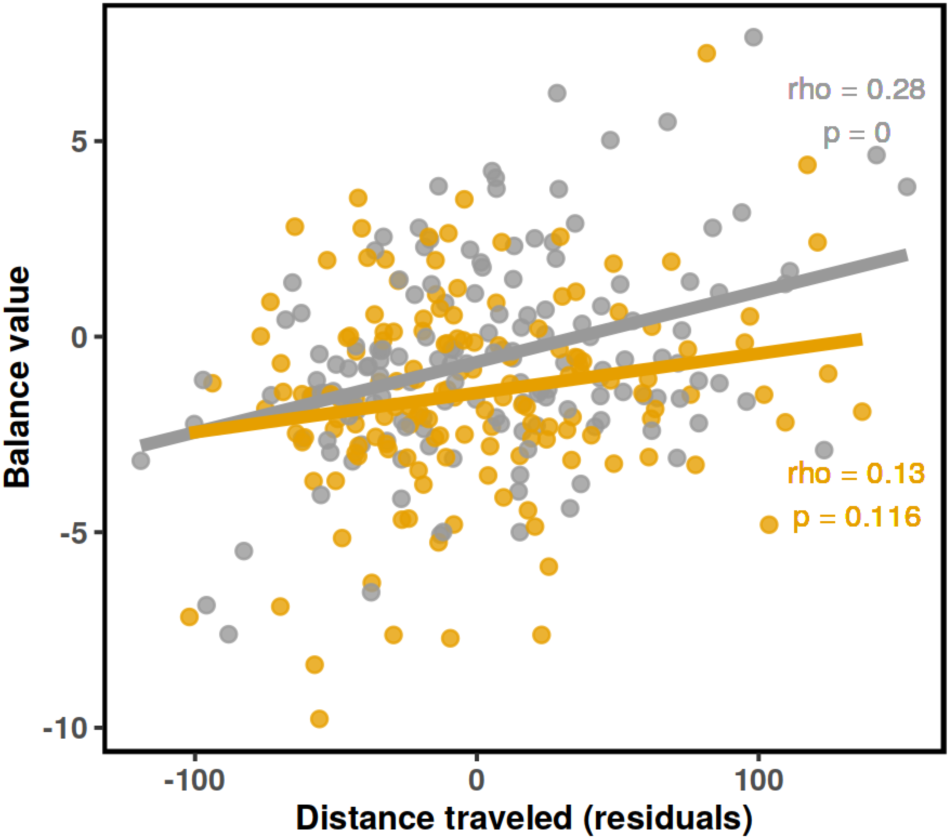
Correlation of the distance traveled adjusted for the body weight at inoculation with the respective nearest balance values, per treatment.

**Figure S7.**
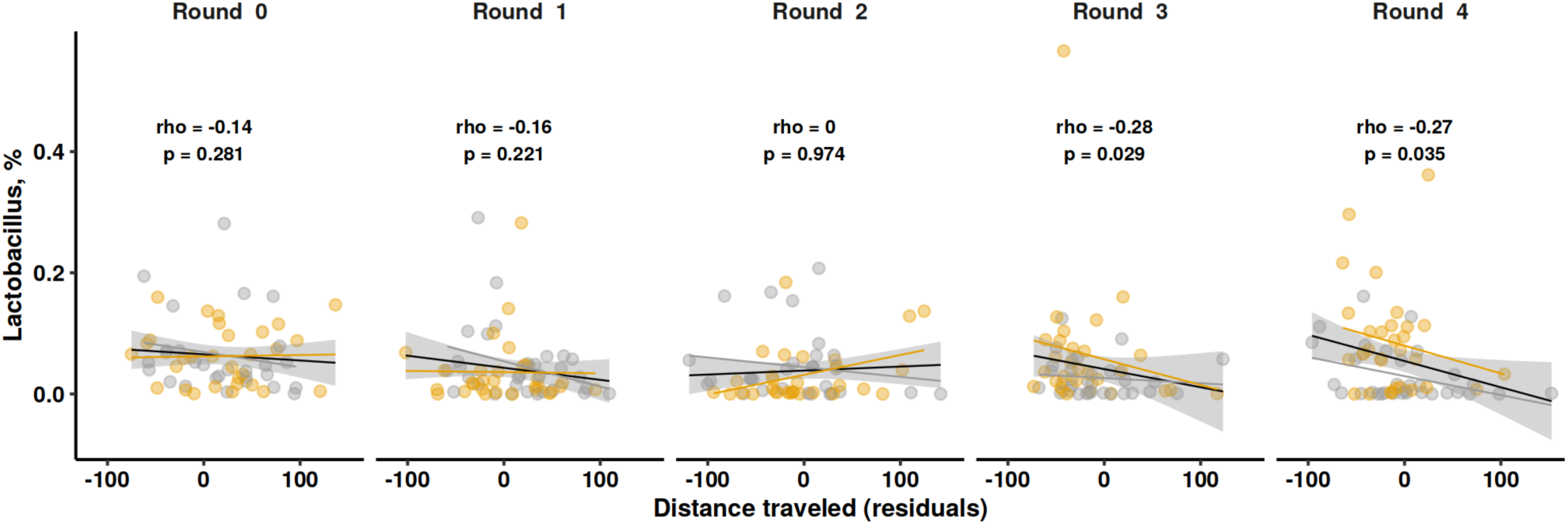
Correlations between distance traveled adjusted for the body weight at inoculation and relative abundance of *Lactobacillus* across the rounds of transfer. Colors indicate the Selection (yellow) and Control lines (gray). The black line shows the linear fit of all data points. Spearman rho and p-value for all data points are shown.

**Figure S8.**
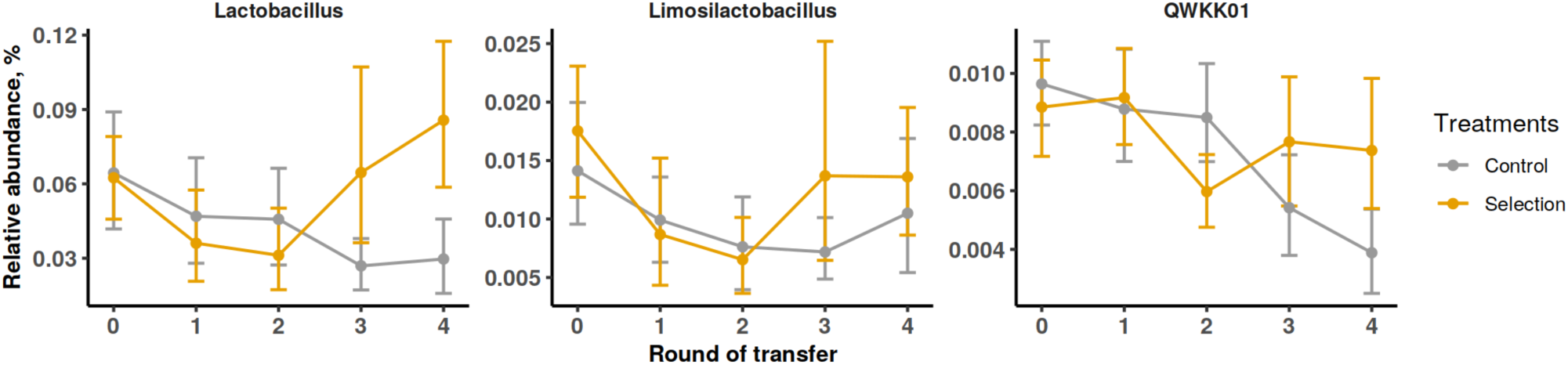
Abundance dynamics of the genera negatively associated with distance traveled. The taxa from the denominator of the respective nearest balance are shown; relative abundance values are clr-transformed values. The first panel is from Fig. 3H for comparison purposes.

**Figure S9.**
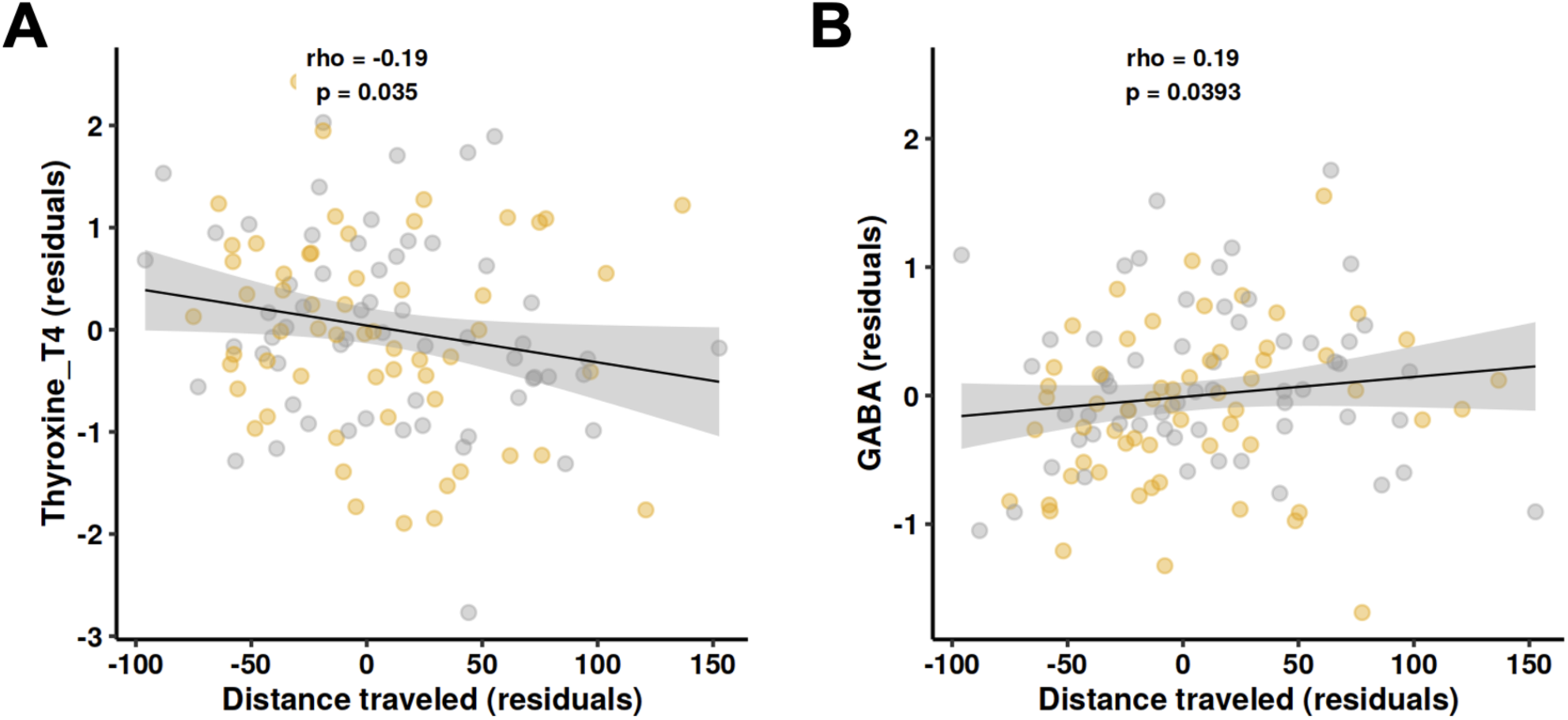
Metabolites significantly correlated with distance traveled. (A) Thyroxine and (B) GABA show significant correlations with distance traveled. Spearman’s rho and uncorrected p-values are shown (n = 118 samples). A significant negative correlation between distance traveled and Indolelactic acid (ILA) is presented in Fig. 4C. Colors indicate the Selection lines (yellow) and Control lines (gray).

**Figure S10.**
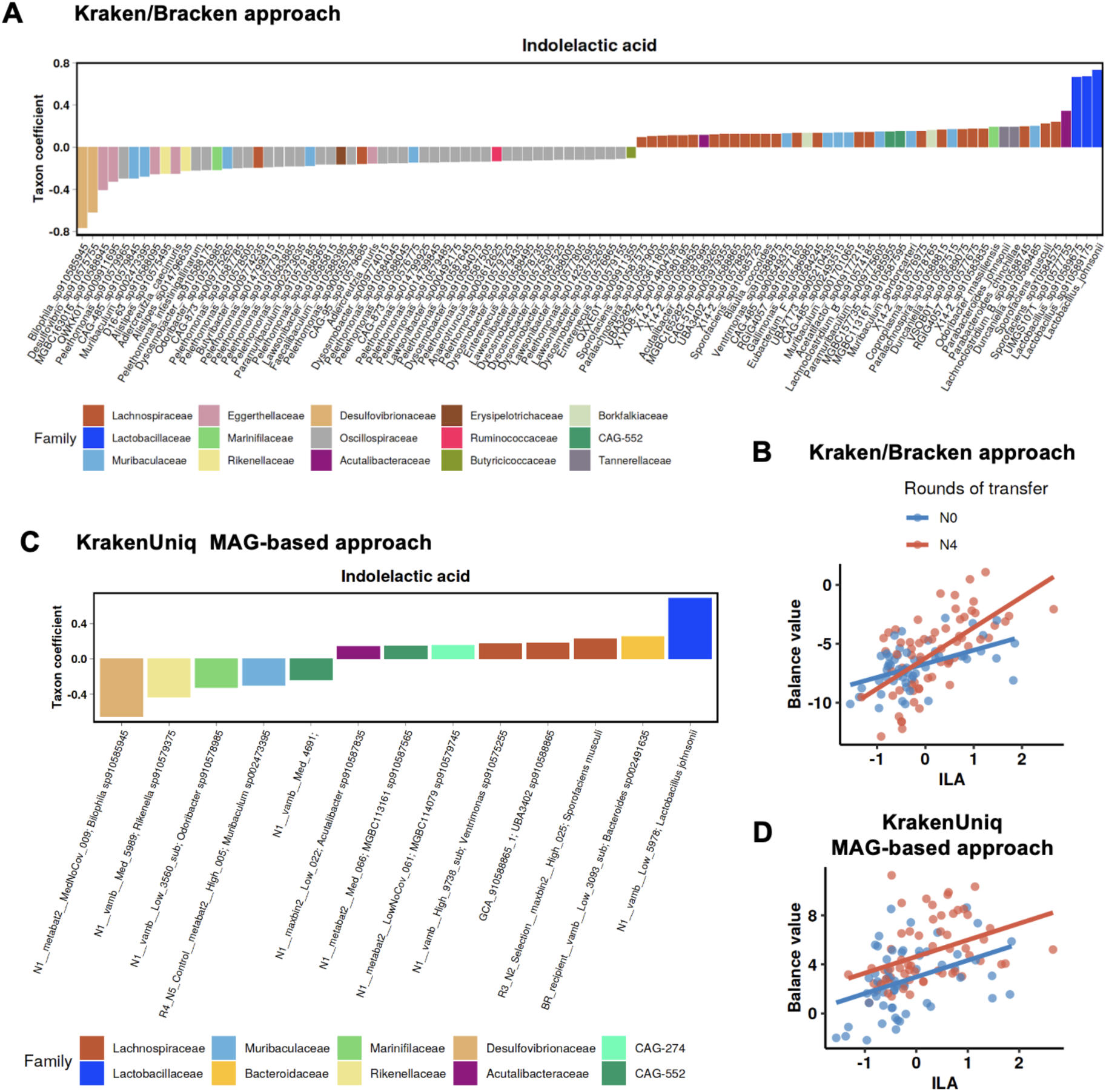
Species-level nearest balance analysis for indolelactic acid (ILA) reveals similar patterns using two approaches. The ILA levels had been adjusted for the body weight at inoculation. A) Based on the default taxonomic profiling method (Kraken/Bracken): species-components of the balance, along with its values (B) calculated for each metagenome with a metabolome available (n=118), with the regression lines per round of transfer. C) Based on the MAGs-based taxonomic profiling (KrakenUniq): species-representative genomes (SRG)-components of the balance (including MAGs’ internal IDs), along with their values (D).

